# Peripheral CB1R inhibition modulates food intake and metabolic efficiency in obesity independently of the gut-brain vagal axis

**DOI:** 10.64898/2025.12.10.693401

**Authors:** Oriane Onimus, Camille de Almeida, Benoit Bertrand, Julien Castel, Nejmeh Mashhour, Anthony Ansoult, Serge Luquet, Giuseppe Gangarossa

## Abstract

**Background and Purpose:** Obesity involves profound disruptions in neuronal circuits, neuroendocrine communication and the endocannabinoid system (ECS). While global cannabinoid type-1 receptor (CB1R) blockade improves metabolism, its clinical use is limited by neuropsychiatric side effects. Peripherally restricted CB1R antagonists offer a safer alternative, yet the neural pathways, specifically the role of the gut-brain vagal axis, mediating their effects remain unclear.

**Experimental Approach:** We investigated the metabolic and neural effects of peripheral inhibition of CB1R (JD5037 and AM6545) in lean and diet-induced obese (DIO). Metabolic parameters were assessed using indirect calorimetry, and neuronal activation was mapped by cFos immunoreactivity. The requirement for vagal signaling was examined using subdiaphragmatic vagotomy (SDV) and pharmacological blockade of cholecystokinin (CCK) and glucagon-like peptide-1 (GLP-1) receptors.

**Key Results:** Peripheral CB1R inhibition suppressed food intake and shifted nutrient partitioning toward fatty acid oxidation in DIO, but not lean, mice. Obesity upregulated CB1R expression in the nodose ganglia. In DIO mice, peripheral CB1R inhibition robustly activated satiety-related brainstem (NTS, AP, PBN) and hypothalamic (ARC, PVN) nuclei. SDV abolished brainstem activation but failed to blunt hypothalamic recruitment or the anorexigenic and metabolic benefits. Furthermore, antagonism of CCK or GLP-1 receptors did not prevent the feeding-suppressive effects of JD5037.

**Conclusions and Implications:** Our findings reveal a dual-mechanism model: vagal pathways mediate brainstem engagement, while hypothalamic recruitment and metabolic improvements occur via vagal-independent signaling. These results demonstrate that peripherally restricted CB1R antagonists indirectly engage central homeostatic circuits, supporting their therapeutic potential for obesity even in conditions with impaired vagal signaling.

**Bullet point summary:** *‘What is already known’:*

- Obesity is associated with elevated circulating endocannabinoids, reflecting chronic overactivation of the peripheral endocannabinoid system.
- Clinical and preclinical studies indicate that peripheral CB1R inhibition ameliorates obesity-related dysfunctions.

*‘What this study adds’:*

- Peripheral CB1R inhibition suppresses food intake and promotes fatty acid oxidation in obese but not lean conditions.
- Peripheral CB1R inhibition engages brainstem nuclei via vagal signaling and hypothalamic nuclei via vagal-independent mechanisms.

*‘Clinical significance’:*

- Metabolic state-dependent CB1R responsiveness must be considered when designing endocannabinoids-targeted obesity therapies.
- Peripheral CB1R antagonists engage central homeostatic circuits via body-brain pathways, warranting surveillance of CNS outcomes.

## Introduction

Obesity is a multifactorial metabolic disease characterized by a chronic positive energy balance, altered eating patterns, excessive fat accumulation, and systemic metabolic dysfunctions. It is associated with increased risks of type 2 diabetes, cardiovascular disease, non-alcoholic fatty liver disease, and several cancers. Growing evidence suggests that obesity may be initiated and/or worsened by distorted functioning of neuronal ensembles as well as of altered body-brain communications [1,2]. However, despite intense research efforts, long-term non-invasive and efficient therapies remain limited, primarily due to the complexity of the underlying regulatory networks that govern energy homeostasis. Among these networks, the endocannabinoid system (ECS) has emerged as a key modulator of both central and peripheral neuronal mechanisms that control appetite, energy storage, glucose metabolism, and lipid homeostasis [3,4].

The ECS consists of endogenous cannabinoids [*i.e.,* <u>anandamide</u> (AEA) and <u>2-arachidonoylglycerol</u> (2-AG)], their receptors [<u>cannabinoid receptor type 1</u> (CB1R) and <u>type</u> <u>2</u> (CB2R)] and the enzymes responsible for their synthesis and degradation. While the role of centrally expressed CB1R in regulating feeding behaviour and hedonic aspects of food intake is well-established [3–5], there is a growing recognition that CB1 receptors located in peripheral tissues (*i.e.,* adipose tissue, liver, muscles, peripheral nervous system and gastrointestinal tract) also contribute to metabolic dysfunctions and represent promising anti-obesity targets [4,6]. In particular, CB1R activation in peripheral organs promotes lipogenesis, impairs insulin sensitivity, and alters gastrointestinal motility and secretion, whereas CB1R inhibition counteracts such effects [7,8]. This therapeutic potential of leveraging the ECS is supported by both clinical and preclinical studies demonstrating elevated levels of AEA and/or 2-AG in several eating disorders [9–12] and also in obesity [13–15], therefore suggesting a constant overactivation of the ECS system in these conditions.

Early pharmacological approaches targeting the ECS, such as the use of <u>rimonabant</u>, a centrally and peripherally active CB1R inverse agonist, demonstrated robust metabolic improvements and weight loss in obese individuals [16]. However, these benefits were overshadowed by significant neuropsychiatric side effects, including anxiety and depression [17–19], due to CB1R antagonism in the brain. This led to the development of peripherally restricted CB1R antagonists and inverse agonists that do not cross the blood-brain barrier [7,8,20,21]. Preclinical studies using such compounds have shown promising effects, including reduction in body weight, improved glucose tolerance, enhanced thermogenesis, and decreased hepatic steatosis, all without eliciting central side effects [7,8,20–25].

An area of active investigation is the extent to which the metabolic and behavioural benefits of peripheral CB1R inhibition are mediated by specific organs [26–29] and also the gut-brain axis [12,30], particularly via the vagal signaling. The vagus nerve serves as a crucial bidirectional conduit between the gut and the brain, transmitting nutritional, hormonal, and microbial signals to central circuits that regulate appetite and metabolism. Peripheral CB1Rs are expressed along the gut epithelium [31,32] as well as in afferent/sensory vagal neurons [12,30,33,34] raising the possibility that pharmacological modulation of CB1R may recruit vagal-dependent mechanisms to influence systemic metabolism.

However, whether the metabolic improvements observed with peripheral CB1R inhibition in obesity are entirely dependent on an intact gut-brain vagal pathway remains unclear. Distinguishing vagus nerve-dependent from vagus nerve-independent mechanisms is essential for optimizing the therapeutic potential of peripheral CB1R inhibition and understanding its full range of physiological and pharmacological effects. Furthermore, clarification of these mechanisms could broaden the scope of CB1R-targeted therapies to patient populations with altered or impaired vagal signaling.

In this study, we investigated whether and how peripheral CB1R inhibition influences energy metabolism under both physiological and pathological conditions, such as obesity. Our findings demonstrate that: (*i*) peripheral CB1R inhibition modulates energy metabolism in obese, but not lean, states; (*ii*) this modulation involves activation of brainstem and hypothalamic nuclei through both vagus-dependent and vagus-independent mechanisms respectively; and (*iii*) critically, the beneficial metabolic effects persist even in the absence of intact gut-brain vagal signaling. These results advance our understanding of endocannabinoid system modulation in metabolic diseases and support the need for continued development of peripherally restricted CB1R-targeted therapies as potentially safe and effective treatments for obesity and eating disorders.

## Material and methods

### Animals

All experimental procedures were approved by the Animal Care Committee of the Université Paris Cité (CEB-22-2019, APAFiS #24407; CEB-38-2021, APAFiS #35447), and carried out following the 2010/63/EU directive. 8-12 weeks old C57BL/6J male mice (Janvier, France) were used and housed in a room maintained at 22 ±1 °C, with a light period from 7h00 to 19h00. Regular chow diet (3.24 kcal/g, reference SAFE® A04, Augy, France), high-fat diet (HFD, Research Diets, Cat #D12492, 5.24 kcal/g) and water were provided *ad libitum* unless otherwise stated. Diet-induced obesity was established by feeding mice a HFD for 2-3 months. All procedures were designed to minimize animal suffering and reduce the number of animals used. For each experiment, mice were randomly assigned to the different treatment groups. Study designs aimed to generate groups of equal or closely matched sizes. Whenever possible, experimenters were blinded to treatment allocation and experimental conditions. Animal studies are reported in compliance with the ARRIVE guidelines [35] and with the recommendations made by the *British Journal of Pharmacology* [36].

### Subdiaphragmatic vagotomy (SDV)

Prior to surgery and during 4-5 post-surgery days, animals were provided with *ad libitum* jelly food (DietGel Boost #72-04-5022, Clear H2O). Animals received Buprécare® (<u>buprenorphine</u> 0.3 mg/kg) and Ketofen® (<u>ketoprofen</u> 10 mg/kg), and were anaesthetized with <u>isoflurane</u> (3.5% for induction, 1.5% for maintenance). During surgery the body temperature was maintained at 37 °C using a heated pad. Briefly, using a binocular microscope, the right and left subdiaphragmatic branches of the vagus nerve were carefully isolated along the lower esophagus/stomach and carefully sectioned in vagotomized animals (SDV mice) or left intact in Sham animals. Mice recovered for at least 3-4 weeks before being used for experimental procedures. The efficiency of the SDV procedure was evaluated as previously described [12,37,38]. Briefly, SDV success was confirmed either prior to or following experiments. SDV mice were included based on at least one of the following criteria: (*i*) lower anorexigenic response to CCK-8S 3-weeks after the SDV/Sham procedure, (*ii*) increased stomach distension, (*iii*) reduced retrograde Fluorogold staining in the dorsal motor nucleus of the vagus (DMV) and/or (*iv*) body weight trajectory during the 3-weeks of post-surgery recovery period [38].

### Drugs

Mice were administered with the following drugs at doses selected from previously published studies: <u>JD5037</u> (3, 6, 10 mg/kg [21,39,40]; #HY-18697, CliniSciences), <u>AM6545</u> (3 mg/kg [41]; #5443, Tocris), <u>devazepide</u> (0.3 mg/kg [31]; #2304, Tocris), <u>exendin-(9-39)</u> (0.1 mg/kg [42,43]; #2081, Tocris), <u>nadolol</u> (10 mg/kg [44]; #HY-B0804, CliniSciences), <u>sotalol</u> hydrochloride (3 mg/kg [45]; #0952, Tocris), <u>glycopyrrolate</u> (0.5 mg/kg [46]; #SML0029, Sigma-Aldrich) and respective vehicle solutions. JD5037 and AM6545 were dissolved in a solution containing DMSO, Kolliphor and saline at the following ratio 2:1:97. Nadolol and glycopyrrolate were dissolved in a solution containing DMSO, PEG300, Tween80 and saline at the following ratio 10:40:5:45. Devazepide, exendin-(9-39) and sotalol were dissolved in saline. Matched vehicles were administered. Devazepide, exendin-(9-39), nadolol and glycopyrrolate were administered 30 minutes prior to JD5037. All drugs were injected intraperitoneally (i.p.) in 10 mL/kg of body volume.

### Tissue preparation and immunofluorescence

Mice were anaesthetized with <u>pentobarbital</u> (500 mg/kg, Dolethal, Vetoquinol, France) and transcardially perfused with cold (4 °C) PFA 4% for 5 min. Brains were post-fixed in PFA 4% at 4°C for 24h and changed in PBS 1X. 40 μm coronal sections were processed using a vibratome (Leica). Confocal imaging acquisitions were performed after immunohistochemistry protocol, using a confocal microscope (Zeiss LSM 710) as previously described [47,48]. The following primary antibodies were used: rabbit anti-cFos (1:1000, Synaptic Systems, #226 003, RRID: AB_2231974, or 1:1000, Cell Signaling, #2250, RRID: AB_2247211). Sections were incubated for 60 min with the following secondary antibodies: donkey anti-rabbit Cy3 AffiniPure (1:1000, Jackson Immunoresearch, 711-165-152, RRID: AB_2307443). Structures were selected according to the following coordinates (from bregma, in mm): PBN (-5.02 to -5.34), AP/NTS (-7.32 to -7.76), PVN (-0.82 to -1.06) and ARC (-1.46 to -1.94). The objectives (10X or 20X) and the pinhole setting (1 airy unit) remained unchanged during the acquisition of a series for all images. Immunopositive cells were quantified using the Cell Counter plugin in ImageJ. A fixed fluorescence threshold was applied as a reference standard. Image acquisition and analysis were performed by experimenters blinded to treatment groups.

### Quantitative RT-PCR

The nodose ganglia were dissected and snap-frozen using liquid nitrogen. All tissues were kept at -80 °C until RNA extraction. Tissues were homogenized in TRIzol/QIAzol Lysis Reagent (Life Technologies) with 3 mm tungsten carbide beads by using the Tissue Lyser III (QIAGEN, 9003240). Total RNA was extracted using the Rneasy Micro Kit (QIAGEN, 74004). The RNA was quantified by using the NanoDrop 1000 spectrophotometer. 500 ng of mRNA from each sample was used for retrotranscription, performed with the SuperScript®III Reverse Transcriptase (Life Technologies) following the manufacturer’s instructions. Quantitative RT-PCRs were performed in a LightCycler 1.5 detection system (Roche, Meylan France) using the Takyon No Rox SYBR MasterMix dTTP Blue (Eurogentec) in 384-well plates according to the manufacturer’s instruction. The following primers were used: *Cnr1* (forward 5’-3’ AGCAAGGACCTGAGACATG, reverse 5’-3’: TGTTATTGGCGTGCTTGTGC), *Rpl19* (forward 5’-3’: GGGCAGGCATATGGGCATA; reverse 5’-3’: GGCGGTCAATCTTCTTGGATT). Relative concentrations were extrapolated from the concentration range for each gene. Concentration values were normalized to the house-keeping gene RPL19.

### Metabolic efficiency analysis

Indirect calorimetry was performed as previously described [12,49]. Mice were monitored for whole energy expenditure (EE), O_2_ consumption, CO_2_ production, respiratory exchange ratio (RER=VCO_2_/VO_2_, V=volume), fatty acid oxidation (FAO) and locomotor activity using calorimetric cages (Labmaster, TSE Systems GmbH, Bad Homburg, Germany). Ratio of gases was determined through an indirect open circuit calorimeter. This system monitors O_2_ and CO_2_ at the inlet ports of a tide cage through which a known flow of air is ventilated (0.4 L/min) and regularly compared to a reference empty cage. O_2_ and CO_2_ were recorded every 15 min during the entire experiment. EE was calculated using the Weir equation for respiratory gas exchange measurements. Food intake was measured with sensitive sensors for automated online measurements. Mice were monitored for body weight and composition at the entry and exit of the experiment using an EchoMRI (Whole Body Composition Analyzers, EchoMRI, Houston, USA). The TSE PhenoMaster software v7.4.2 was used to collect and visualize the data. Data were initially processed in Microsoft Excel using extracted raw values of VO_2_, VCO_2_ (mL/15 min), and energy expenditure (EE; kcal/15 min). Statistical analyses were subsequently performed using GraphPad Prism 10 (GraphPad Software, La Jolla, CA, USA). Covariance analyses between energy expenditure (EE) and lean body mass (LBM) are reported in **Suppl. Fig. 1** and **2**.

### Transcriptomics meta-analysis

Publicly available transcriptomic data [50] were downloaded from Gene Expression Omnibus (https://www.ncbi.nlm.nih.gov/geo/, GSE138651) and analysed using a Python 3.0 pipeline generated in line with the original publication.

### Statistics

All data are presented as mean ± SEM. Statistical tests were performed with Prism 10 (GraphPad Software, La Jolla, CA, USA). No a priori sample size calculation was performed. Statistical analysis was undertaken only for studies where each group size was at least n=5. The declared group size is the number of independent values, and statistical analysis was done using these independent values. Sample sizes (n = 5-12 mice per experimental group) were chosen (*i*) based on previous studies using similar methodologies and in line with standard practice in the field, and (*ii*) to comply with ethical considerations aimed at minimizing animal use, in accordance with the 3Rs principle, without compromising data quality. Normal distribution of data was analysed using the Shapiro-Wilk test. Depending on the experimental design, data were analysed using either Student’s t-test with equal variances, One-way ANOVA or Two-way ANOVA. The significance threshold was automatically set at p<0.05. ANOVA analyses were followed by Bonferroni *post hoc* test for specific comparisons only when overall ANOVA revealed a significant difference (at least p<0.05).

### Nomenclature of Targets and Ligands

Key protein targets and ligands in this article are hyperlinked to corresponding entries in http://www.guidetopharmacology.org, the common portal for data from the IUPHAR/BPS Guide to Pharmacology, and are permanently archived in the Concise Guide to PHARMACOLOGY 2023/24 [51].

## Results

### The metabolic efficacy of peripheral CB1R inhibition depends on the metabolic state

Recent studies have shown that pharmacological inhibition of peripheral CB1 receptors using agents such as JD5037 or AM6545 holds strong promise for treating metabolic and eating disorders (for review see [52]). Here, we explored whether and how peripheral CB1R inhibition affects feeding behaviour and metabolic efficiency. To address this, we first administered different doses of the peripherally restricted CB1R inverse agonist JD5037 (3, 6, and 10 mg/kg, i.p.) to lean mice fed a standard chow diet (CD) and assessed various metabolic parameters of energy homeostasis. Interestingly, JD5037 had no significant effect on food intake (**Fig. 1A, E, I**), respiratory exchange ratio (**Fig. 1B, F, J**), fatty acid oxidation (**Fig. 1C, G, K**), or energy expenditure (**Fig. 1D, H, L**), compared to vehicle-treated controls. This set of results indicate that inhibition of peripheral CB1R in lean animals does not alter feeding, nutrient partitioning (oxidation of carbohydrates *versus* lipids) or energy balance.

**Figure 1:**
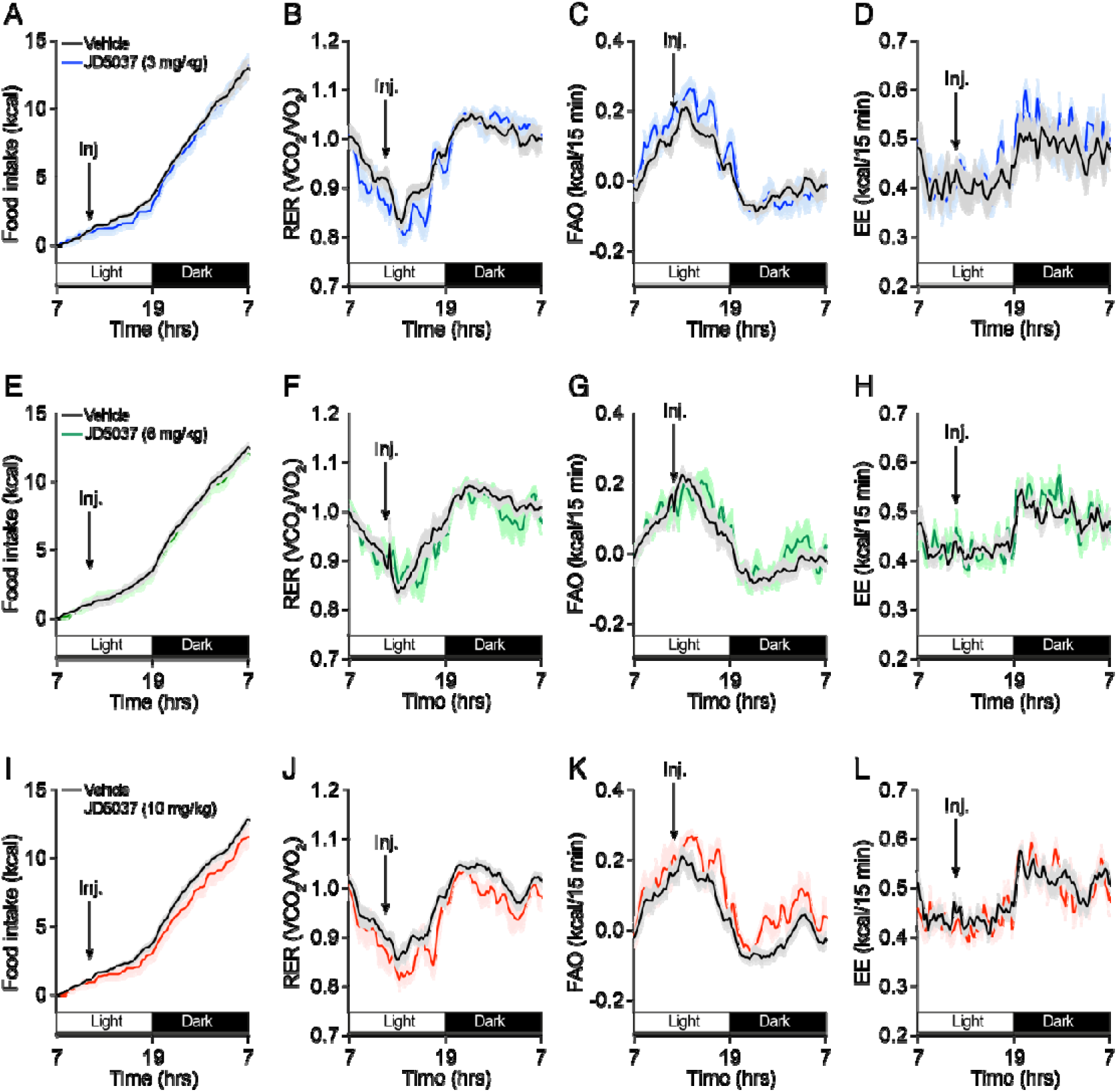
Inhibition of peripheral CB1R does not affect feeding and energy homeostasis in lean mice. Escalating doses (3, 6, 10 mg/kg, i.p.) of the peripheral CB1R inverse agonist JD5037 were administered to lean mice, and their feeding and metabolic profiles were characterized by using metabolic chambers (indirect calorimetry). (**A, E, I**) Food intake, (**B, F, J**) respiratory exchange ratio (RER), (**C, G, K**) fatty acid oxidation (FAO) and (**D, H, L**) energy expenditure (EE) in mice injected with vehicle or JD5037 (3, 6, 10 mg/kg) (n=6-7 mice/group). Statistics: Two-way ANOVA followed by a Bonferroni *post hoc* test (all panels), not significant.

Next, we investigated whether the beneficial effects of peripheral CB1R inhibition depended on the metabolic state. Mice were fed a high-fat diet (HFD) for 2-3 months, leading to a significant increase in body weight (**Fig. 2A**). In contrast to lean mice, administration of the peripheral CB1R inverse agonist JD5037 (3 mg/kg, i.p.) markedly reduced food intake in HFD-fed mice (**Fig. 2B, B^1^**). This was accompanied by a shift in nutrient utilization favouring lipids over carbohydrates, as evidenced by a decreased respiratory exchange ratio (RER; **Fig. 2C, C^1^**) and an increased fatty acid oxidation (FAO; **Fig. 2D, D^1^**). These metabolic changes occurred without significant alterations in energy expenditure (EE; **Fig. 2E, E^1^, Suppl. Fig. 1A**) or spontaneous locomotor activity (**Fig. 2F**). Beside the metabolic state (chow diet-*versus* HFD-mice), we also wondered whether the nutritional state (fasted *versus* fed mice) may condition the anorexigenic effect of JD5037. Again, we observed that JD5037 (3 mg/kg, i.p.) maintained its anorexigenic properties only in HFD-mice (food consumed after 4 hours of refeeding, **Fig. 2G**).

**Figure 2:**
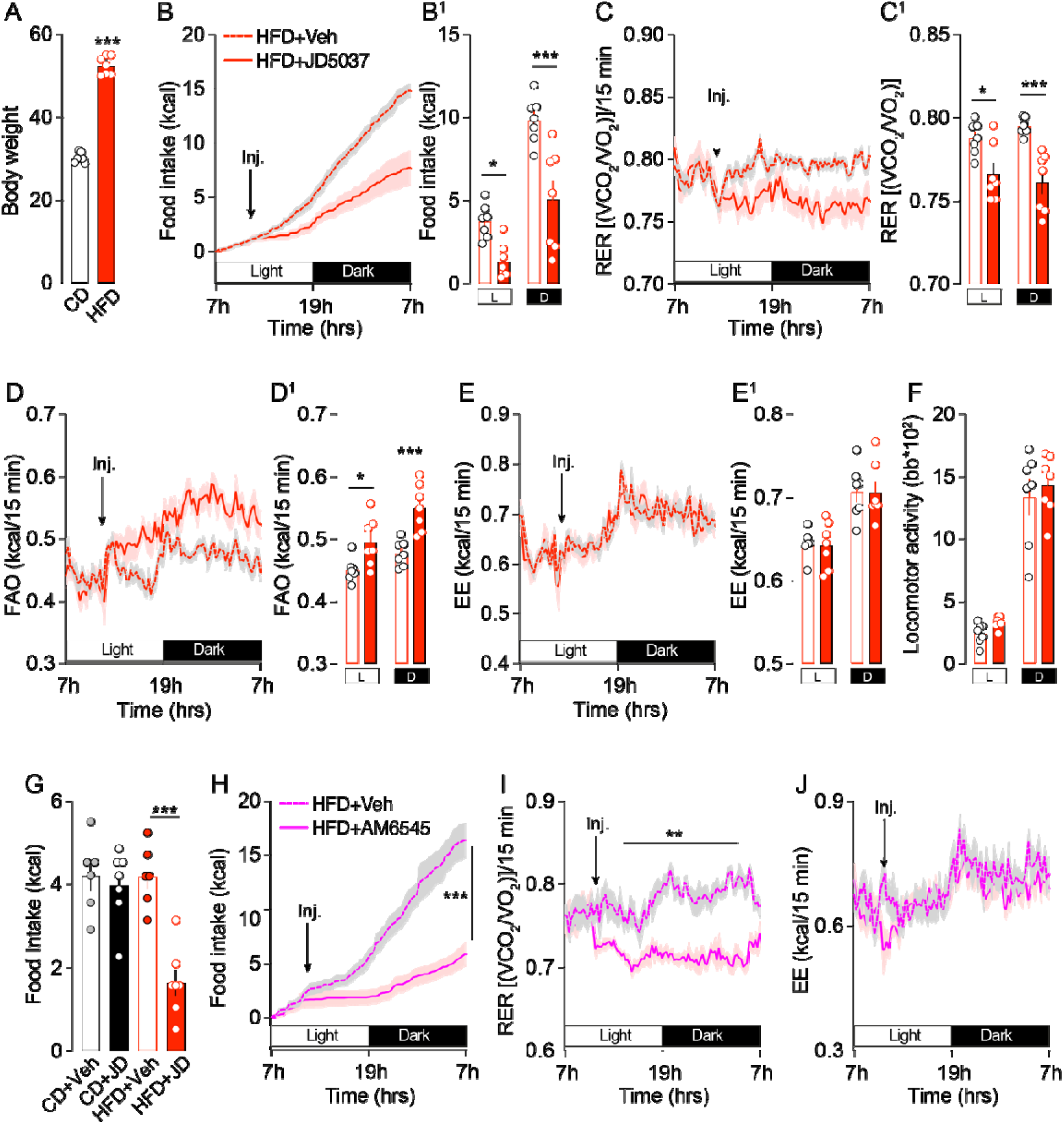
Inhibition of peripheral CB1R affects feeding and nutrient partitioning in HFD-mice. (**A**) HFD-induced body weight gain. Veh or JD5037 (3 mg/kg, i.p.) were administered to HFD-mice and their feeding and metabolic profiles were characterized by using metabolic chambers (indirect calorimetry). (**B, B^1^**) Food intake, (**C, C^1^**) respiratory exchange ratio (RER), (**D, D^1^**) fatty acid oxidation (FAO), (**E, E^1^,** see also **Suppl. Fig. 1A**) energy expenditure (EE) and (**F**) spontaneous locomotor activity in HFD-mice injected with vehicle or JD5037 (3 mg/kg, i.p.) (n=7 mice/group). (**G**) Food intake in fasted chow diet- and HFD-mice 4 hours after refeeding schedule (n=6-7 mice/group). (**H**) Food intake, (**I**) RER and (**J,** see also **Suppl. Fig. 1B**) EE in HFD-mice injected with vehicle or the peripheral CB1R neutral antagonist AM6545 (3 mg/kg, i.p.) (n=7 mice/group). Statistics: Student’s t-test (A panel), Two-way ANOVA followed by a Bonferroni *post hoc* test (B^1^, C^1^, D^1^, E^1^, F, H, I, J panels), One-way ANOVA followed by a Bonferroni *post hoc* test (G panel), *p<0.05, **p<0.01 and ***p<0.001 for specific comparisons.

To further complement these findings, we used the peripherally restricted CB1R antagonist AM6545 (3 mg/kg, i.p.) and observed comparable outcomes: a reduction in food intake (**Fig. 2H**) and a shift in nutrient partitioning (**Fig. 2I**), again without affecting energy expenditure (**Fig. 2J, Suppl. Fig. 1B**).

### Inhibition of peripheral CB1R activates satietogenic brainstem nuclei in a metabolic state-dependent manner via the gut-brain vagal axis

Building on our findings related to feeding behaviour and energy metabolism, we next examined whether peripheral CB1R inhibition triggers specific patterns of neuronal activation in key brainstem nuclei involved in satiety regulation, namely the nucleus tractus solitarius (NTS), area postrema (AP), and parabrachial nucleus (PBN) (**Fig. 3A**), which are known to participate in anorexigenic/satietogenic responses [53–56].

**Figure 3:**
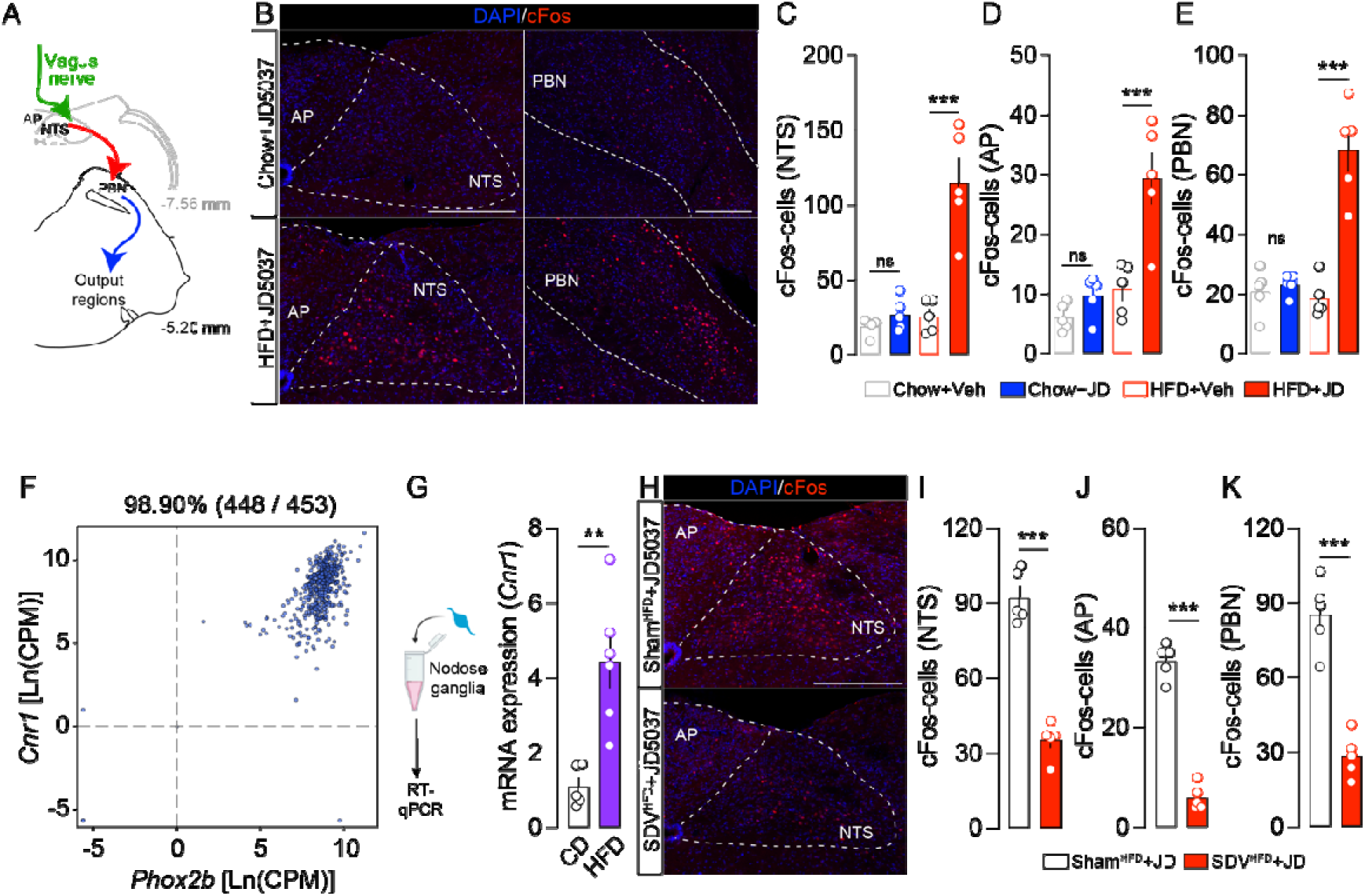
Inhibition of peripheral CB1R leads to the activation of brainstem nuclei via a vagus nerve-dependent mechanism. (**A**) The drawing illustrates vagus nerve-mediated activation of brainstem nuclei. (**B**) Immunofluorescence detection of cFos in the nucleus tractus solitarius (NTS), area postrema (AP), and parabrachial nucleus (PBN) of chow- and HFD-mice administered with JD5037 (3 mg/kg, i.p.). Quantifications of cFos-positive cells in the NTS (**C**), AP (**D**) and PBN (**E**) of chow- and HFD-mice administered with vehicle or JD5037 (3 mg/kg) (n=5 mice/group). (**F**) Transcriptomic meta-analysis of *Phox2b* and *Cnr1* in sensory neurons (n=453) of the nodose ganglia (NG). (**G**) Relative mRNA expression of *Cnr1* in the nodose ganglia of chow- and HFD-mice (n=5-6 mice/group). (**H**) Immunofluorescence detection and quantifications (**I, J, K**) of cFos-positive cells in the NTS, AP and PBN of Sham^HFD^- and SDV^HFD^-mice administered with JD5037 (3 mg/kg, i.p.) (n=5 mice/group). Scale bars: 250 μm. Statistics: One-way ANOVA followed by a Bonferroni *post hoc* test (C, D, E panels) or Student’s t-test (G, I, J, K panels), **p<0.01 and ***p<0.001 for specific comparisons.

Chow- and HFD-fed mice were treated with JD5037 (3 mg/kg, i.p.) or vehicle, and neuronal activity was assessed in the NTS, AP, and PBN using cFos immunoreactivity as a *bonafide* molecular proxy of neuronal activation. Strikingly, and in line with our earlier metabolic observations (**Fig. 1**, **2**), JD5037 increased the number of cFos-positive cells in these brainstem regions exclusively in HFD-fed mice (**Fig. 3A-E**). This suggests that peripheral CB1R inhibition engages periphery-to-brain satiety circuits in a metabolic state-dependent manner.

Eating disorders and obesity are associated with a chronic elevation of circulating endocannabinoids, particularly 2-arachidonoylglycerol (2-AG) and/or anandamide (AEA), in both humans and animal models, reflecting a systemic overactivation of the endocannabinoid system (ECS) [11,15,57–59]. Given the activation of the NTS observed in HFD-fed mice following JD5037 administration, we hypothesized that the vagus nerve, which conveys peripheral metabolic/nutritional signals to NTS-neurons, undergoes CB1R-related molecular adaptations. To explore this aspect, we first examined whether neurons in the nodose ganglia (NG) express CB1R. Consistent with previous reports [12,33], transcriptomic meta-analysis [50] revealed a strong enrichment of *Cnr1* (the transcript encoding for CB1R) in *Phox2b*-positive neurons of the NG with ∼99% of vagal neurons also co-expressing *Cnr1* (**Fig. 3F**). Furthermore, direct quantification of *Cnr1* expression in the NG showed a significant upregulation in HFD-fed mice compared to chow-fed controls (**Fig. 3G**), suggesting that obesity sensitizes vagal afferents to endocannabinoid signaling.

To directly assess whether JD5037-induced cFos activation in the NTS/AP was mediated via the gut-brain vagal axis, we employed the subdiaphragmatic vagotomy (SDV) model. In HFD-fed mice that underwent SDV (SDV^HFD^-mice), JD5037 failed to elicit the robust cFos activation observed in Sham^HFD^-mice, with a dramatic reduction of cFos expression in the NTS, AP and PBN (**Fig. 3H-K**). These findings indicate that peripheral CB1R inhibition activates interconnected satiety-associated brainstem nuclei through a vagus nerve-dependent mechanism.

### The gut-brain vagal axis is dispensable for the metabolic effects of peripheral CB1R inhibition

Although the gut-brain vagal axis is required for JD5037-induced neuronal activation in the brainstem, we next wondered whether this pathway was also necessary for the metabolic effects of peripheral CB1R inhibition. To this end, we compared JD5037-mediated metabolic responses in Sham^HFD^- *versus* SDV^HFD^-mice by using calorimetric cages. In this experiment, food was removed during the light phase (**Fig. 4A, B**; magenta area) to isolate metabolic responses from food intake (**Fig. 4A, B**; green area).

**Figure 4:**
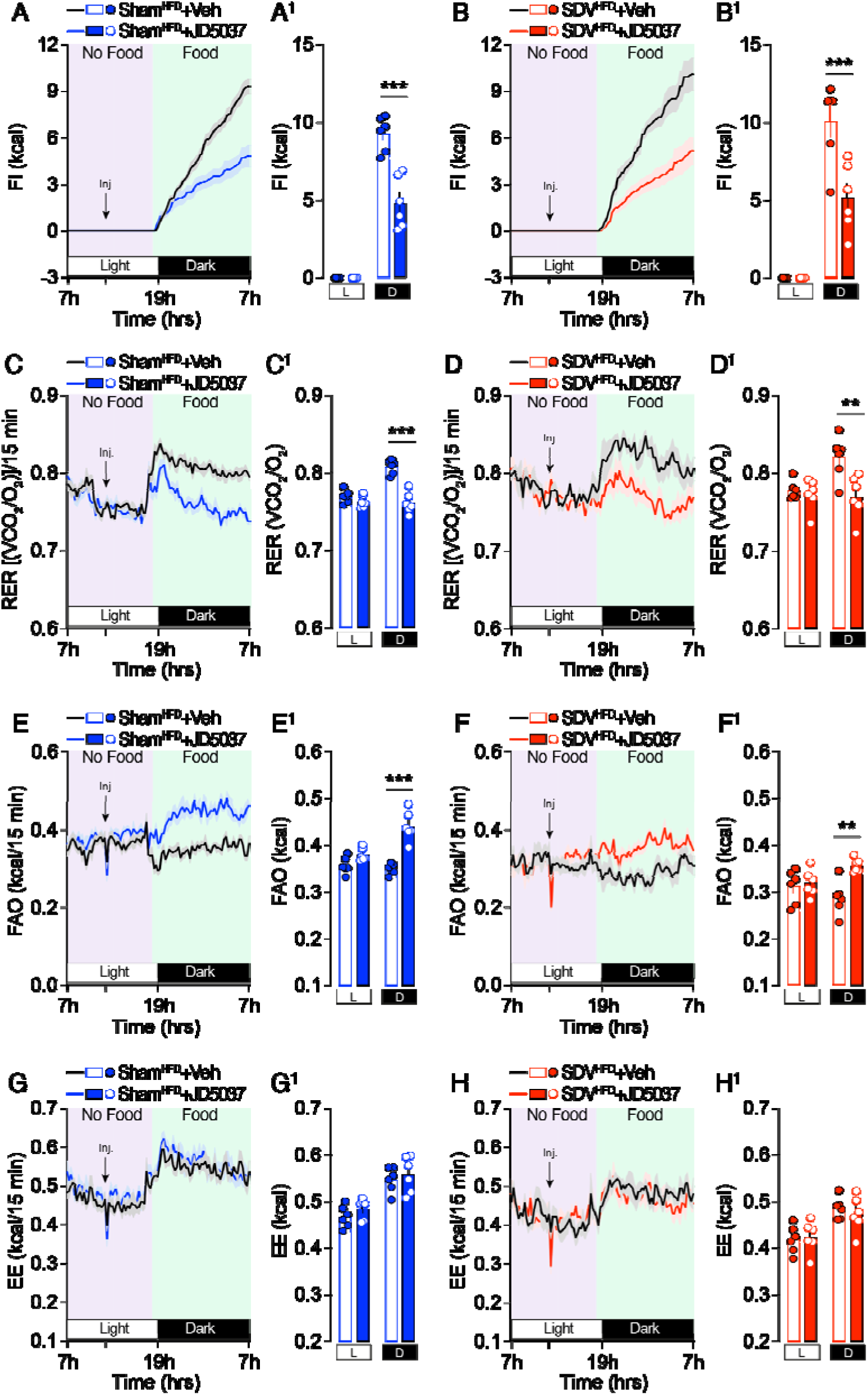
The gut-brain vagal axis does not mediate JD5037-induced feeding and metabolic adaptations. Veh or JD5037 (3 mg/kg, i.p.) were administered to Sham^HFD^- and SDV^HFD^-mice and their feeding and metabolic profiles were characterized by using metabolic chambers (indirect calorimetry). (**A, A^1^, B, B^1^**) Food intake, (**C, C^1^, D, D^1^**) respiratory exchange ratio (RER), (**E, E^1^, F, F^1^**) fatty acid oxidation (FAO) and (**G, G^1^, H, H^1^,** see also **Suppl. Fig. 2**) energy expenditure in Sham^HFD^- and SDV^HFD^-mice injected with vehicle or JD5037 (3 mg/kg) (n=6 mice/group). Statistics: Two-way ANOVA followed by a Bonferroni *post hoc* test (all panels), **p<0.01 and ***p<0.001 for specific comparisons.

During this fasting period, JD5037 administration had no impact on respiratory exchange ratio (RER), fatty acid oxidation (FAO), or energy expenditure (EE) in either group (**Fig. 4C-H**), confirming that food intake is required to reveal the metabolic shift of nutrient partitioning and utilization. When food was reintroduced prior to the dark phase (**Fig. 4A, B**; green area), JD5037 significantly reduced food intake in Sham^HFD^-mice (**Fig. 4A, A^1^**) but surprisingly also in SDV^HFD^-mice (**Fig. 4B, B^1^**). In addition, JD5037-induced changes in nutrient utilization, characterized by a reduced RER and an enhanced FAO, were observed in both groups (**Fig. 4C-F^1^**), with no significant differences in energy expenditure (**Fig. 4G-H^1^, Suppl. Fig. 2**). These results suggest that, although the gut-brain vagal axis is essential for the central activation of satiety-associated brainstem nuclei (**Fig. 3**), it is not required for either the metabolic effects or the changes in food intake (FI) induced by peripheral CB1R inhibition.

To further validate the dispensability of gut-brain vagal signaling in the metabolic effects of peripheral CB1R inhibition, we pharmacologically blocked the action of two key gut-derived hormones: cholecystokinin (CCK) and glucagon-like peptide-1 (GLP-1). These incretins are (*i*) known to be regulated by CB1R [27,31,60,61] and (*ii*) well-established activators of vagal afferents [62,63]. We co-administered the <u>CCK receptor</u> antagonist devazepide (**Fig. 5A**) or the <u>GLP-1 receptor</u> antagonist exendin-(9-39) (**Fig. 5B**) along with JD5037 in both chow- and HFD-fed mice. Strikingly, neither antagonist reversed the anorexigenic effect of JD5037 (**Fig. 5A, B**), indicating that blocking CCK or GLP-1 signaling does not impair the appetite-suppressing action of peripheral CB1R inhibition. These findings further support the conclusion that gut-brain endocrine signaling is not required for the metabolic benefits elicited by peripherally acting CB1R antagonists in a HFD model.

**Figure 5:**
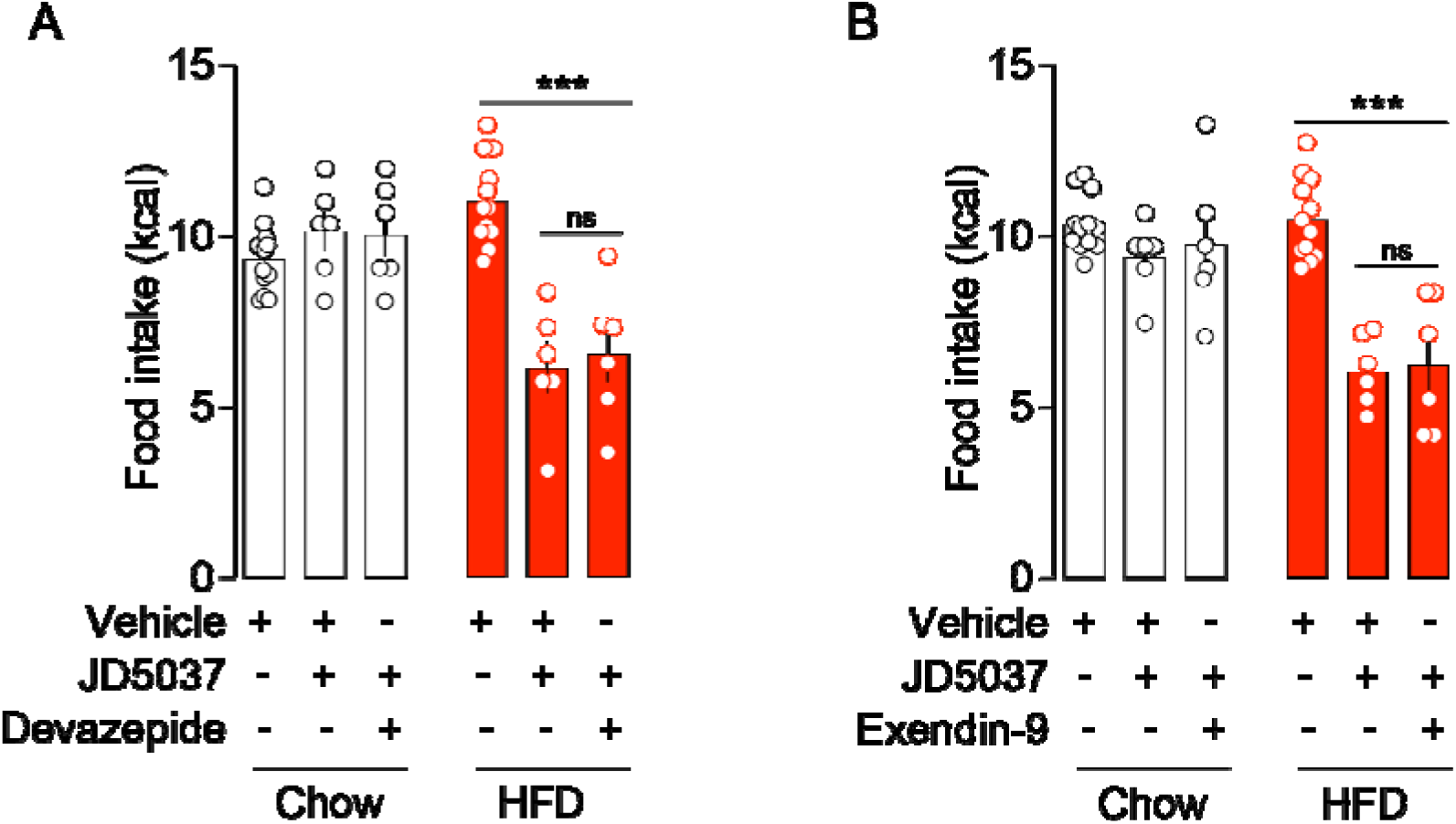
CCK and GLP-1 do not mediate JD5037-induced feeding adaptations. (**A, B**) Food intake in lean and HFD-mice following administration the CCK receptor antagonist devazepide (0.3 mg/kg) (**A**) or the GLP1 receptor antagonist Exendin-9 (0.1 mg/kg) (**B**) (n=6-12 mice/group). Statistics: Two-way ANOVA followed by a Bonferroni *post hoc* test (all panels), ***p<0.001 for specific comparisons.

### Inhibition of peripheral CB1R activates satietogenic hypothalamic nuclei in a metabolic state-dependent but a vagus nerve-independent manner

The hypothalamus is a central regulator of feeding behaviour and energy homeostasis [64]. We therefore examined whether peripheral CB1R inhibition triggered activation of two major hypothalamic nuclei involved in appetite regulation: the paraventricular nucleus (PVN) and the arcuate nucleus (ARC). JD5037 significantly increased the number of cFos-positive cells in both regions, but only in HFD-fed mice (**Fig. 6A-C**), reinforcing our earlier observations in the brainstem (**Fig. 3**) that peripheral CB1R inhibition engages periphery-to-brain satiety pathways in a metabolic state-dependent manner.

**Figure 6:**
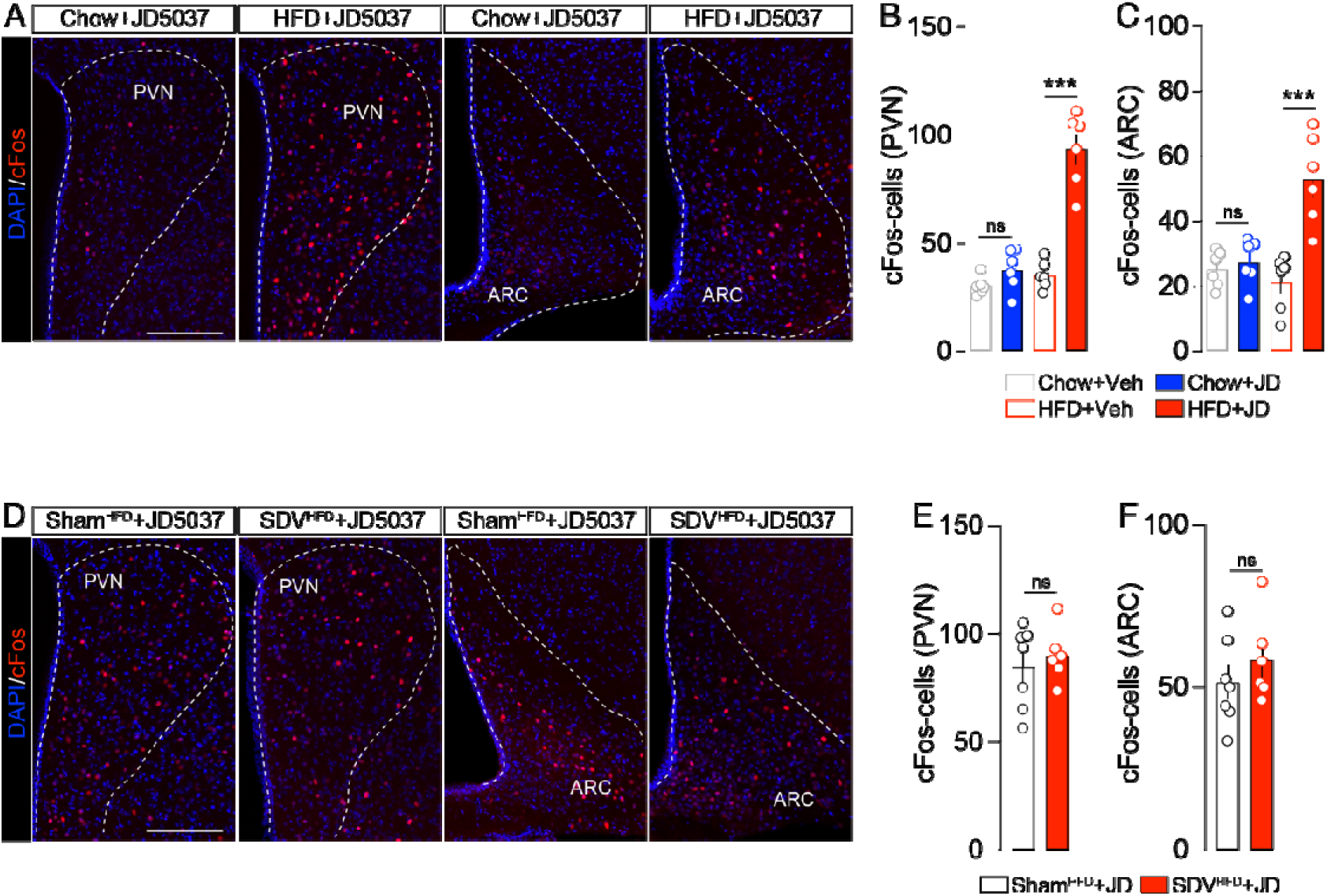
Inhibition of peripheral CB1R leads to the activation of hypothalamic nuclei via a vagus nerve-independent mechanism. (**A**) Immunofluorescence detection of cFos in the paraventricular nucleus (PVN) and arcuate nucleus (ARC) of the hypothalamus of chow-and HFD-mice administered with JD5037 (3 mg/kg). Quantifications of cFos-positive cells in the PVN (**B**) and ARC (**C**) of chow- and HFD-mice administered with vehicle or JD5037 (3 mg/kg) (n=6 mice/group). (**D**) Immunofluorescence detection and quantifications (**E, F**) of cFos-positive cells in the PVN and ARC of Sham^HFD^- and SDV^HFD^-mice administered with JD5037 (3 mg/kg) (n=6-7 mice/group). Scale bars: 150 μm. Statistics: One-way ANOVA followed by a Bonferroni *post hoc* test (B, C panels) or Student’s t-test (E, F panels), ***p<0.001 for specific comparisons.

We next investigated whether this hypothalamic activation required an intact vagus nerve signaling. Unexpectedly, JD5037 still induced robust activation of both PVN- and ARC-neurons in Sham^HFD^- and SDV^HFD^-mice (**Fig. 6D-F**). Thus, in contrast to the brainstem (**Fig. 3H-K**), hypothalamic recruitment by peripheral CB1R inhibition appears to occur via a vagus-independent mechanism. This alternative pathway may account for the comparable behavioural and metabolic improvements observed in Sham^HFD^- and SDV^HFD^-mice (**Fig. 4**), supporting the existence of a distinct body-brain communication route driving hypothalamic engagement.

To further explore the mechanisms underlying this effect, we assessed the potential involvement of the autonomic nervous system. In fact, previous studies have shown that the hypophagic action of rimonabant, a brain-penetrant CB1R antagonist, depends on the activation of the sympathetic nervous system [45,65]. In contrast, pre-treatment of HFD-mice with the peripherally restricted sympathetic blockers nadolol (10 mg/kg, i.p.) and sotalol (3 mg/kg, i.p.) did not alter JD5037-induced hypophagia (**Suppl. Fig. 3A, B**). Likewise, inhibition of the parasympathetic tone using the peripheral cholinergic antagonist glycopyrrolate (0.5 mg/kg, i.p.) failed to affect the anorectic response of JD5037 (**Suppl. Fig. 3C**).

Collectively, these findings demonstrate that peripheral CB1R inhibition activates hypothalamic satiety circuits through a mechanism that is independent of both vagal and autonomic nervous system signaling.

## Discussion

Our study provides compelling evidence that the metabolic efficacy of peripheral CB1 receptor (CB1R) inhibition is strongly dependent on the metabolic state, with clear distinctions between lean and diet-induced obese (DIO) mice. In fact, while peripherally restricted CB1R inverse agonists such as JD5037 had no measurable impact on feeding behaviour, nutrient partitioning, or energy expenditure in lean animals, the same treatment induced robust anorexigenic and metabolic effects in HFD-fed mice. These findings underscore the importance of the metabolic environment in modulating responsiveness to peripheral endocannabinoid system (ECS) manipulation and are consistent with clinical and preclinical evidence indicating an exacerbated peripheral endocannabinoid tone in metabolic and eating disorders [3,6].

A key finding is also that inhibition of peripheral CB1R selectively reduced food intake and shifted fuel utilization towards fatty acid oxidation in HFD-fed mice, consistent with previous studies showing that ECS overactivation contributes to hyperphagia and metabolic dysfunctions in obesity [3]. This metabolic state dependence likely reflects (mal)adaptive changes in peripheral ECS tone, as supported by the upregulation of ECS-related signaling components and CB1R in several peripheral tissues [14,57,66,67], as well as in the nodose ganglia of HFD-fed mice (current study). This structure, which houses the sensory neurons of the vagus nerve and integrates peripheral interoceptive and metabolic signals, may be particularly relevant. In fact, increased vagal *Cnr1* expression may contribute, at least in part, to the molecular and functional rearrangements underlying the blunted vagal activity induced by chronic exposure to obesogenic diets [68–71].

Interestingly, we observed that peripheral CB1R inhibition activated satiety-associated brainstem circuits, specifically the NTS, AP, and PBN, but only in HFD-fed animals. This activation, visualized via increased cFos immunoreactivity and most likely resulting from a CB1R-coupled G_i_-dependent disinhibition, required an intact gut-brain vagal axis, as subdiaphragmatic vagotomy strongly blunted JD5037-induced cFos expression in the brainstem. These data suggest that obesity-associated ECS overactivity sensitizes the gut-brain vagal axis to peripheral CB1R inhibition, thereby engaging central networks.

However, our findings also reveal a crucial dissociation: the gut-brain vagal axis is necessary for neuronal activation in the brainstem but is dispensable for the metabolic effects of peripheral CB1R inhibition. In fact, even in SDV^HFD^-mice, JD5037 retained its ability to reduce food intake and shift nutrient partitioning, suggesting that peripheral mechanisms, independent of gut-brain vagal inputs, are sufficient to mediate these homeostatic/metabolic outcomes. These results, which are in line with a previous study showing that CB1R in vagal *Phox2b*-neurons is dispensable for HFD-induced body weight gain [30], highlight that the vagal endocannabinoid tone is mobilized in specific physio-pathological contexts such as binge eating [12], alcohol preference/consumption [72,73], refeeding in fasted states [74] but not obesity, which is a particular metabolic disorder. Furthermore, here we demonstrated that pharmacological blockade of the gut hormones CCK and GLP-1, whose release and activity are scaled by CB1R-related signaling [27,31,60,75], did not prevent the anorexigenic effects of JD5037. This finding further reinforces the notion that gut-derived endocrine signals are not critical mediators of the metabolic effects elicited by peripheral CB1R inhibition. Thus, the persistence of JD5037 efficacy in the presence of CCK and GLP-1 antagonism suggests alternative, non-vagal mechanisms that may be responsible for the metabolic improvements observed. These non-vagal, CB1R-mediated body-brain mechanisms may involve the release and/or clearance of mediators from CB1R-expressing peripheral organs [13,28,76–79] in the bloodstream and/or the recruitment of sensory spinal afferents [80,81], thereby modulating feeding and metabolic outputs independently of gut-brain vagal inputs.

This interpretation is reinforced by our findings showing that the hypothalamic PVN and ARC nuclei of obese mice, similar to brainstem nuclei, exhibit hypersensitivity to peripheral CB1R inhibition, but through a mechanism independent of the vagus nerve. Importantly, this observation contrasts with our previous report demonstrating that peripheral CB1R inhibition induces a vagus-dependent activation of PVN- and ARC-neurons in binge-eating mice [12], an effect that is absent in obese animals in the present study. This apparent discrepancy in vagal involvement across metabolic states may reflect state-dependent alterations of the endocannabinoid system. Indeed, while binge-eating mice are characterized by elevated levels of 2-AG without significant changes in AEA [12], obese individuals exhibit changes in both endocannabinoids [14,82]. Such differences in ligand availability may critically influence how peripheral signals are relayed to the brain. Consistent with this idea, recent evidence indicates that 2-AG and AEA can differentially engage CB1R depending on receptor localization and cell types, at least in the hippocampus, thereby eliciting distinct effects on synaptic transmission and plasticity [83]. Together, these findings suggest that ligand-specific and context-dependent CB1R signaling may underlie the differential recruitment of vagal pathways across metabolic states.

In this context, our results support the existence of alternative body-brain communication pathways capable of engaging hypothalamic circuits independently of the gut-brain vagal axis. Recently, chronic administration of JD5037 has been shown to restore <u>leptin</u> sensitivity in obese mice, while also increasing circulating <u>adiponectin</u> and free fatty acids (FFAs) [7,21]. These parallel adipose tissue-derived factors may cross the blood-brain barrier and converge at the hypothalamic level in a vagus nerve-independent manner, thereby promoting the satiety and metabolic effects associated with JD5037, independently of brainstem nuclei activation. An open question remains regarding the functional role of vagus-dependent brainstem circuits activated (cFos induction) following peripheral CB1R inhibition. One plausible hypothesis is that CB1R-mediated vagal signaling does not primarily influence homeostatic feeding and peripheral energy metabolism, but instead modulates the hedonic, reward-driven aspects of food intake [12,38,84]. Future studies should explore whether the reward deficits observed in obese individuals may, at least in part, stem from dysfunctions in the gut-brain endocannabinoid vagal system. This possibility is supported by evidence that pharmacological elevation of 2-AG levels using the <u>MAGL</u> inhibitor <u>JZL184</u> induces binge-eating behaviour in a vagus-dependent manner [12].

Taken together, our results support a dual-pathway model for the action of peripheral CB1R antagonists in obesity. One mechanistic pathway involves vagus-dependent neuronal activation of brainstem nuclei. The second is a vagus-independent homeostatic and metabolic pathway, likely involving direct peripheral effects on organs such as adipose tissue and/or the liver, which may release endocrine mediators that ultimately act on the hypothalamus. This model reconciles the CNS effects of peripheral CB1R inhibition [12,85–87] with the sustained metabolic benefits seen in the absence of vagal or incretin signaling.

Our findings have important implications for the development of peripherally restricted CB1R antagonists as therapeutic strategies for obesity and related metabolic disorders, particularly in light of recent clinical trials reporting significant metabolic improvements with the peripheral CB1R inhibitor <u>monlunabant</u> [88,89]. The state-dependent responsiveness to peripheral CB1R inhibition highlights the need to consider individual metabolic states when evaluating therapeutic efficacy. Moreover, the dissociation between neuronal activation and metabolic improvements suggests that engagement of satiety-associated brainstem circuits, while desirable, may not be required for metabolic benefits, potentially reducing the risk of CNS-mediated side effects. However, our study also highlights that targeting the peripheral endocannabinoid system is not devoid of central consequences, as peripheral signals can influence the activity of key brain nuclei through body-brain communication pathways. Although peripherally restricted CB1R antagonists are expected to produce fewer adverse effects than brain-penetrant compounds, further investigation is required to fully delineate their effects on metabolic, behavioural, and cognitive functions, as well as on associated patterns of brain activity and circuit dynamics.

Future work should focus on identifying the peripheral cellular targets of CB1R inhibition that mediate the observed shifts in energy balance and nutrient utilization. It will also be important to determine whether similar mechanisms operate in humans and whether biomarkers of ECS overactivity (*e.g.,* circulating endocannabinoid levels or CB1R expression profiles) can predict responsiveness to peripheral CB1R-based therapies.

## Acknowledgments

We thank Olja Kacanski for administrative support; Isabelle Le Parco, Daniel Quintas, Magguy Boa, Ludovic Maingault, Angélique Dauvin and Florianne Michel for animals’ care. We acknowledge the Functional and Physiological Exploration platform (FPE) of the Université Paris Cité, CNRS, *Unité de Biologie Fonctionnelle et Adaptative*, and the animal core facility “Buffon” of the Université Paris Cité/Institut Jacques Monod. We also thank Fabrice Licata of the BioMedTech Facilities (INSERM US36, CNRS UAR2009, Université Paris Cité) for help with data acquisition and technical advice.

## Funding

This work was supported by the Agence Nationale de la Recherche (ANR-21-CE14-0021-01, ANR-23-CE14-0014-02, ANR-24-CE14-1322-03), Fédération pour la Recherche sur le Cerveau (FRC), Institut universtaire de France (IUF), Plan d’investissement France 2030 and Idex Emergence (ANR-18-IdEX-0001), IReSP (AAP-2023-SPA-13), Université Paris Cité and CNRS (to G.G.). O.O. is supported by a PhD fellowship from the Fondation pour la Recherche Médicale (FRM). G.G. was also partially supported by EMBO and the Alexander von Humboldt Foundation.

## Author contributions

Oriane Onimus: Conceptualization; Investigation; Methodology; Visualization; Formal analysis; Data curation; Writing - original draft; Writing - review & editing; Software. Camille de Almeida: Investigation; Methodology; Formal analysis. Benoit Bertrand: Investigation; Methodology; Formal analysis. Julien Castel: Investigation; Methodology. Nejmeh Mashhour: Investigation; Methodology. Anthony Ansoult: Investigation; Methodology. Serge Luquet: Resources. Giuseppe Gangarossa: Conceptualization; Investigation; Funding acquisition; Writing - original draft; Methodology; Validation; Visualization; Writing - review & editing; Formal analysis; Project administration; Software; Data curation; Supervision; Resources.

## Competing interests

The other authors declare no competing interests.

## Data availability

The data that support the findings of this study are available from the corresponding author upon reasonable request.

## Declaration of transparency and scientific rigour

This Declaration acknowledges that this paper adheres to the principles for transparent reporting and scientific rigour of preclinical research as stated in the BJP guidelines for Design and Analysis, Immunoblotting and Immunochemistry, and Animal Experimentation, and as recommended by funding agencies, publishers and other organisations engaged with supporting research.

## Abbreviations

AP: area postrema
ARC: arcuate nucleus of the hypothalamus
CB1R: cannabinoid type-1 receptor
CCK: cholecystokinin
CD: chow diet
DIO: diet-induced obesity
ECS: endocannabinoid system
EE: energy expenditure
FAO: fatty acid oxidation
GLP-1: glucagon-like peptide-1
HFD: high-fat diet
NTS: nucleus tractus solitarius
PBN: parabrachial nucleus
PVN: paraventricular nucleus of the hypothalamus
RER: respiratory exchange ratio

---

**Suppl. Figure 1:**
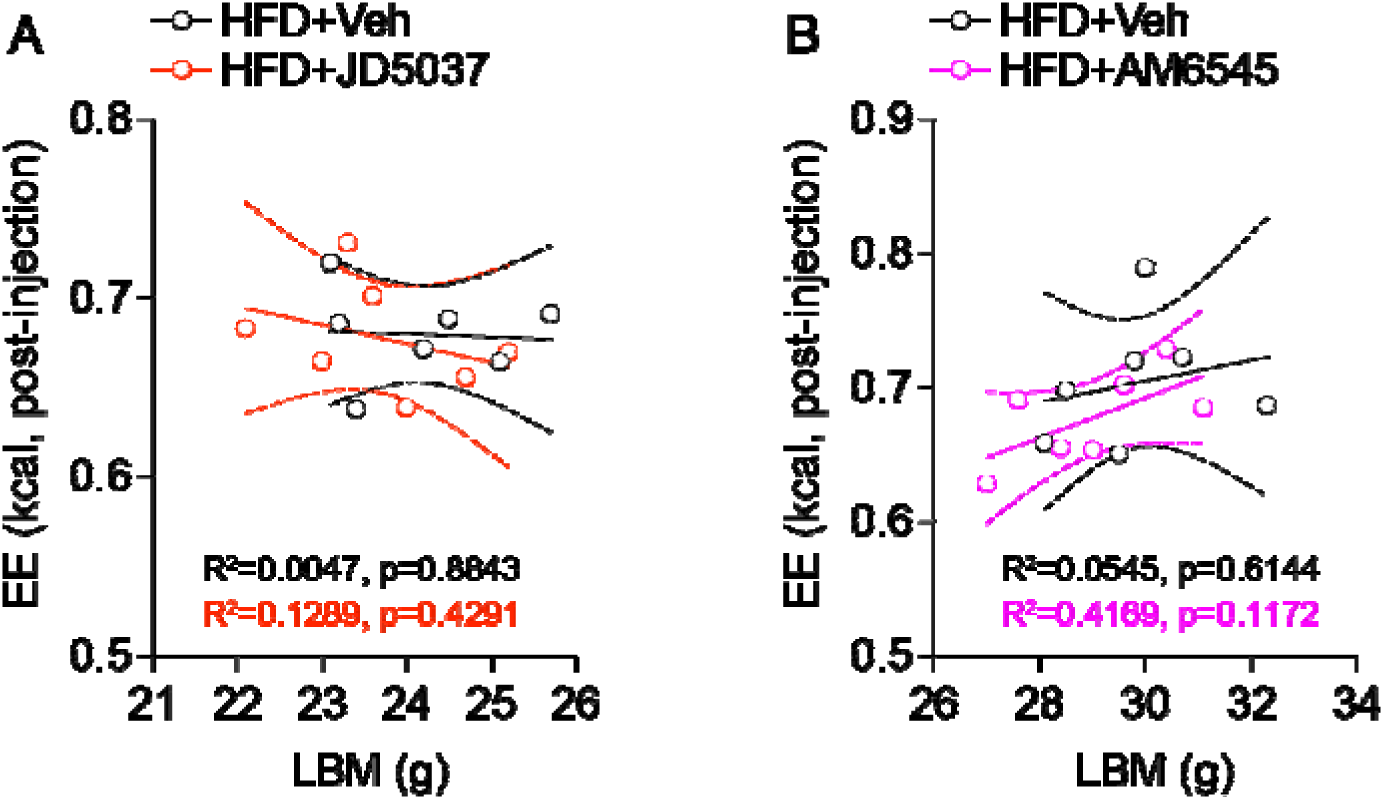
Energy expenditure does not covary with lean body mass in HFD-mice. Analysis of the relationship between energy expenditure (EE) and lean body mass (LBM) in HFD-mice treated with vehicle or JD5037 (**A**), or vehicle or AM6545 (**B**). This analysis relates to Figure 2. Statistics: R^2^ and p values are indicated within the panels.

**Suppl. Figure 2:**
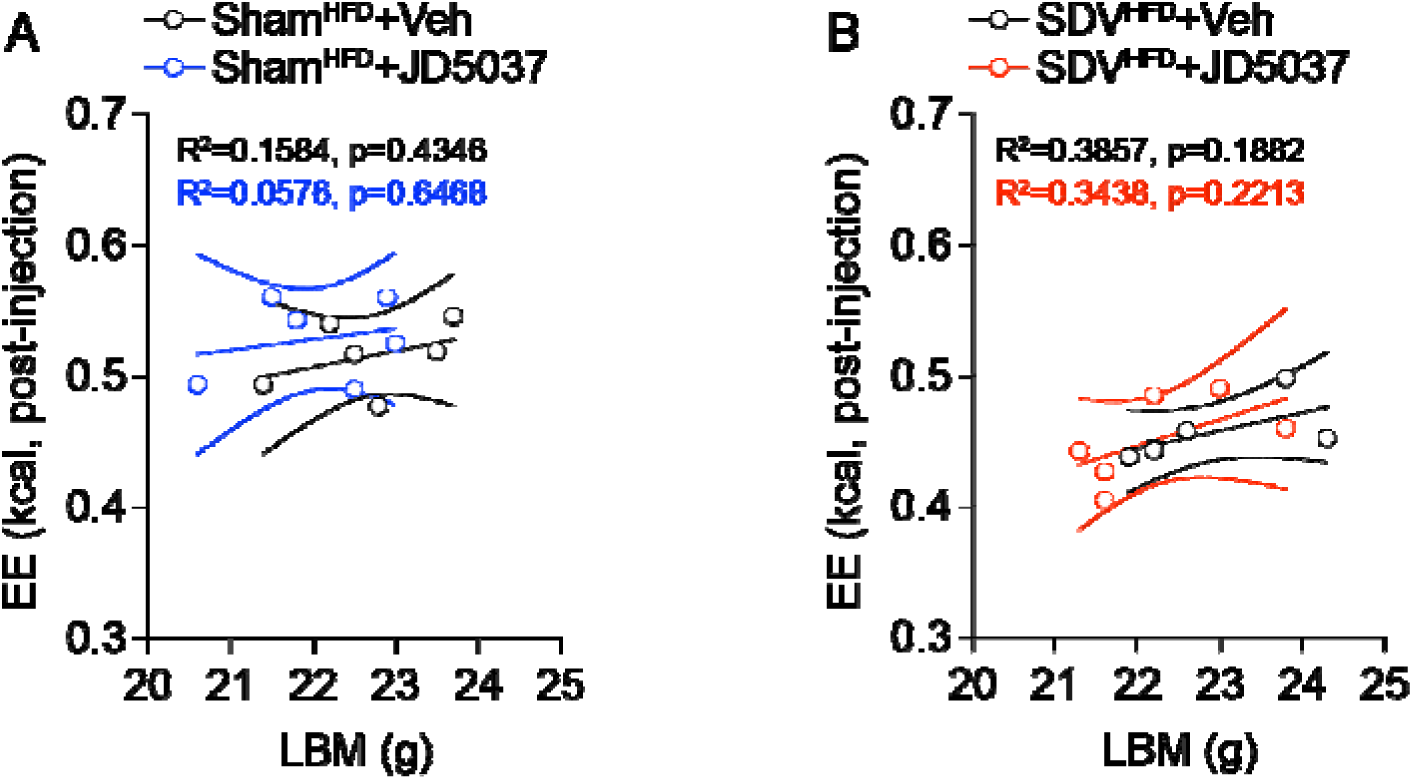
Energy expenditure does not covary with lean body mass in Sham^HFD^-and SDV^HFD^-mice. Analysis of the relationship between energy expenditure (EE) and lean body mass (LBM) in Sham^HFD^- and SDV^HFD^-mice treated with vehicle or JD5037 (**A** for Sham^HFD^-mice; **B** for SDV^HFD^-mice). This analysis relates to Figure 4. Statistics: R^2^ and p values are indicated within the panels.

**Suppl. Figure 3:**
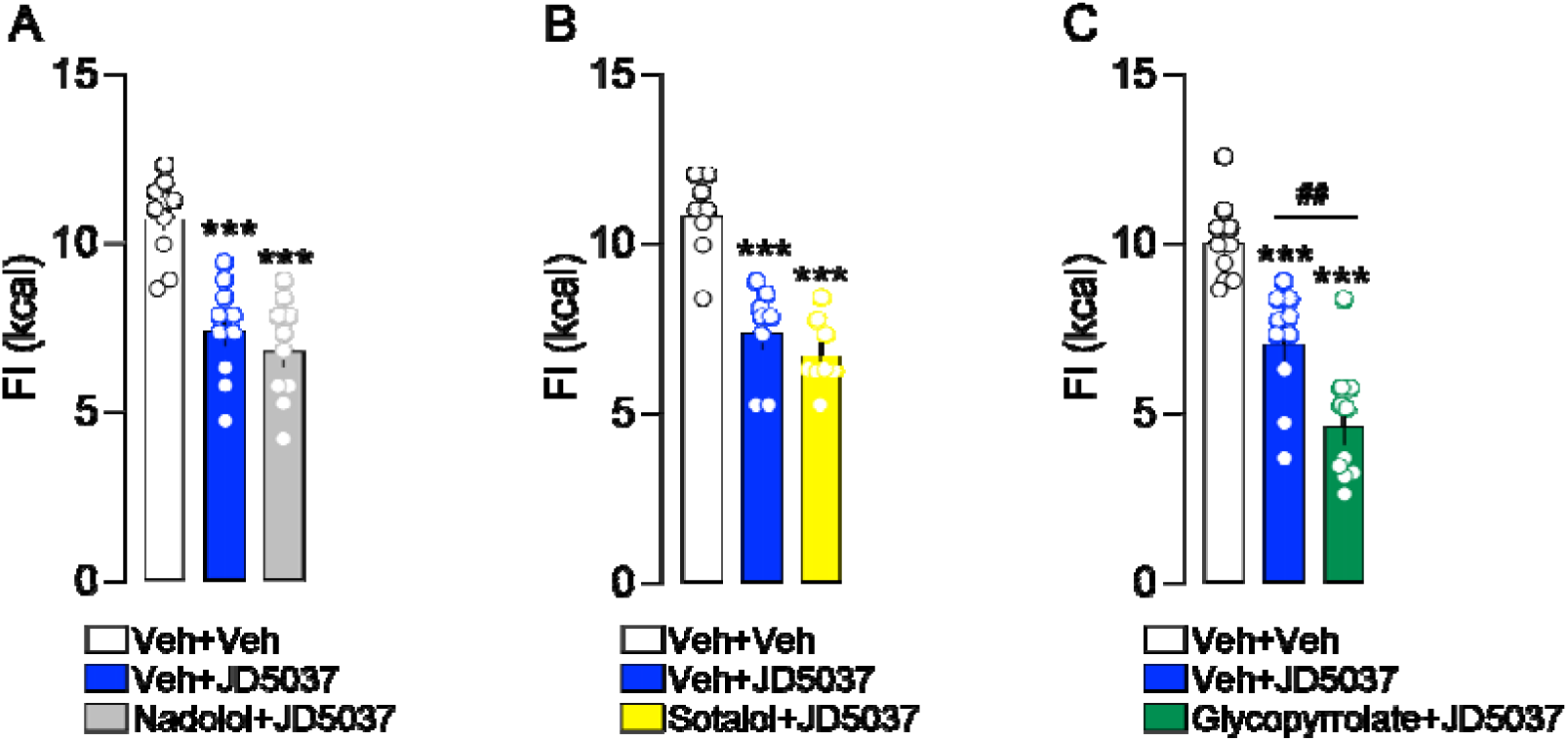
The anorexigenic effect of JD5037 in HFD-mice does not depend on recruitment of the autonomic nervous system. Food intake (FI) following administration of JD5037 in combination with the sympathetic blockers nadolol (**A**, n=10/group) and sotalol (**B**, n=8/group), or the parasympathetic blocker glycopyrrolate (**C**, n=10/group) in HFD-mice. Statistics: One-way ANOVA followed by Bonferroni *post hoc* test (all panels); ***p<0.001 for comparisons with the Veh+Veh group; ^##^p<0.01 for comparisons with the Veh+JD5037 group.

## Notes

### Competing Interest Statement

The authors have declared no competing interest.

### Summary of Updates

New experiments to reinforce our findings.

## References

[1] A. Furlan, P. Petrus, Brain-body communication in metabolic control, Trends Endocrinol Metab 34 (2023) 813–822. 10.1016/j.tem.2023.08.014.

[2] S. Saeed, A. Bonnefond, P. Froguel, Obesity: exploring its connection to brain function through genetic and genomic perspectives, Mol Psychiatry 30 (2025) 651–658. 10.1038/s41380-024-02737-9.

[3] W. Mazier, N. Saucisse, B. Gatta-Cherifi, D. Cota, The Endocannabinoid System: Pivotal Orchestrator of Obesity and Metabolic Disease, Trends Endocrinol Metab 26 (2015) 524–537. 10.1016/j.tem.2015.07.007.

[4] C. Quarta, R. Mazza, S. Obici, R. Pasquali, U. Pagotto, Energy balance regulation by endocannabinoids at central and peripheral levels, Trends Mol Med 17 (2011) 518–526. 10.1016/j.molmed.2011.05.002.

[5] B.K. Lau, D. Cota, L. Cristino, S.L. Borgland, Endocannabinoid modulation of homeostatic and non-homeostatic feeding circuits, Neuropharmacology 124 (2017) 38–51. 10.1016/j.neuropharm.2017.05.033.

[6] C. Quarta, D. Cota, Anti-obesity therapy with peripheral CB1 blockers: from promise to safe(?) practice, Int J Obes (Lond) 44 (2020) 2179–2193. 10.1038/s41366-020-0577-8.

[7] J. Tam, G. Szanda, A. Drori, Z. Liu, R. Cinar, Y. Kashiwaya, M.L. Reitman, G. Kunos, Peripheral cannabinoid-1 receptor blockade restores hypothalamic leptin signaling, Mol Metab 6 (2017) 1113–1125. 10.1016/j.molmet.2017.06.010.

[8] J. Tam, V.K. Vemuri, J. Liu, S. Bátkai, B. Mukhopadhyay, G. Godlewski, D. Osei-Hyiaman, S. Ohnuma, S.V. Ambudkar, J. Pickel, A. Makriyannis, G. Kunos, Peripheral CB1 cannabinoid receptor blockade improves cardiometabolic risk in mouse models of obesity, J Clin Invest 120 (2010) 2953–2966. 10.1172/JCI42551.

[9] I. Knani, B.J. Earley, S. Udi, A. Nemirovski, R. Hadar, A. Gammal, R. Cinar, H.J. Hirsch, Y. Pollak, I. Gross, T. Eldar-Geva, D.P. Reyes-Capo, J.C. Han, A.M. Haqq, V. Gross-Tsur, R. Wevrick, J. Tam, Targeting the endocannabinoid/CB1 receptor system for treating obesity in Prader-Willi syndrome, Mol Metab 5 (2016) 1187–1199. 10.1016/j.molmet.2016.10.004.

[10] A.M. Monteleone, F. Piscitelli, R. Dalle Grave, M. El Ghoch, V. Di Marzo, M. Maj, P. Monteleone, Peripheral Endocannabinoid Responses to Hedonic Eating in Binge-Eating Disorder, Nutrients 9 (2017). 10.3390/nu9121377.

[11] P. Monteleone, I. Matias, V. Martiadis, L. De Petrocellis, M. Maj, V. Di Marzo, Blood levels of the endocannabinoid anandamide are increased in anorexia nervosa and in binge-eating disorder, but not in bulimia nervosa, Neuropsychopharmacology 30 (2005) 1216–1221. 10.1038/sj.npp.1300695.

[12] C. Berland, J. Castel, R. Terrasi, E. Montalban, E. Foppen, C. Martin, G.G. Muccioli, S. Luquet, G. Gangarossa, Identification of an endocannabinoid gut-brain vagal mechanism controlling food reward and energy homeostasis, Mol Psychiatry 27 (2022) 2340–2354. 10.1038/s41380-021-01428-z.

[13] D.A. Argueta, N.V. DiPatrizio, Peripheral endocannabinoid signaling controls hyperphagia in western diet-induced obesity, Physiol Behav 171 (2017) 32–39. 10.1016/j.physbeh.2016.12.044.

[14] E.N. Kuipers, V. Kantae, B.C.E. Maarse, S.M. van den Berg, R. van Eenige, K.J. Nahon, A. Reifel-Miller, T. Coskun, M.P.J. de Winther, E. Lutgens, S. Kooijman, A.C. Harms, T. Hankemeier, M. van der Stelt, P.C.N. Rensen, M.R. Boon, High Fat Diet Increases Circulating Endocannabinoids Accompanied by Increased Synthesis Enzymes in Adipose Tissue, Front Physiol 9 (2018) 1913. 10.3389/fphys.2018.01913.

[15] T.J. Little, N. Cvijanovic, N.V. DiPatrizio, D.A. Argueta, C.K. Rayner, C. Feinle-Bisset, R.L. Young, Plasma endocannabinoid levels in lean, overweight, and obese humans: relationships to intestinal permeability markers, inflammation, and incretin secretion, Am J Physiol Endocrinol Metab 315 (2018) E489–E495. 10.1152/ajpendo.00355.2017.

[16] V. Di Marzo, CB(1) receptor antagonism: biological basis for metabolic effects, Drug Discov Today 13 (2008) 1026–1041. 10.1016/j.drudis.2008.09.001.

[17] C.E. Beyer, J.M. Dwyer, M.J. Piesla, B.J. Platt, R. Shen, Z. Rahman, K. Chan, M.T. Manners, T.A. Samad, J.D. Kennedy, B. Bingham, G.T. Whiteside, Depression-like phenotype following chronic CB1 receptor antagonism, Neurobiol Dis 39 (2010) 148–155. 10.1016/j.nbd.2010.03.020.

[18] D.R. Janero, A. Makriyannis, Cannabinoid receptor antagonists: pharmacological opportunities, clinical experience, and translational prognosis, Expert Opin Emerg Drugs 14 (2009) 43–65. 10.1517/14728210902736568.

[19] B. Le Foll, A. Pushparaj, Y. Pryslawsky, B. Forget, K. Vemuri, A. Makriyannis, J.M. Trigo, Translational strategies for therapeutic development in nicotine addiction: rethinking the conventional bench to bedside approach, Prog Neuropsychopharmacol Biol Psychiatry 52 (2014) 86–93. 10.1016/j.pnpbp.2013.10.009.

[20] N.L. Cluny, V.K. Vemuri, A.P. Chambers, C.L. Limebeer, H. Bedard, J.T. Wood, B. Lutz, A. Zimmer, L.A. Parker, A. Makriyannis, K.A. Sharkey, A novel peripherally restricted cannabinoid receptor antagonist, AM6545, reduces food intake and body weight, but does not cause malaise, in rodents, Br J Pharmacol 161 (2010) 629–642. 10.1111/j.1476-5381.2010.00908.x.

[21] J. Tam, R. Cinar, J. Liu, G. Godlewski, D. Wesley, T. Jourdan, G. Szanda, B. Mukhopadhyay, L. Chedester, J.-S. Liow, R.B. Innis, K. Cheng, K.C. Rice, J.R. Deschamps, R.J. Chorvat, J.F. McElroy, G. Kunos, Peripheral cannabinoid-1 receptor inverse agonism reduces obesity by reversing leptin resistance, Cell Metab 16 (2012) 167–179. 10.1016/j.cmet.2012.07.002.

[22] J. Liu, G. Godlewski, T. Jourdan, Z. Liu, R. Cinar, K. Xiong, G. Kunos, Cannabinoid-1 Receptor Antagonism Improves Glycemic Control and Increases Energy Expenditure Through Sirtuin-1/Mechanistic Target of Rapamycin Complex 2 and 5’Adenosine Monophosphate-Activated Protein Kinase Signaling, Hepatology 69 (2019) 1535–1548. 10.1002/hep.30364.

[23] C. Roger, C. Buch, T. Muller, J. Leemput, L. Demizieux, P. Passilly-Degrace, R. Cinar, M.R. Iyer, G. Kunos, B. Vergès, P. Degrace, T. Jourdan, Simultaneous Inhibition of Peripheral CB1R and iNOS Mitigates Obesity-Related Dyslipidemia Through Distinct Mechanisms, Diabetes 69 (2020) 2120–2132. 10.2337/db20-0078.

[24] S. Tan, H. Liu, B. Ke, J. Jiang, B. Wu, The peripheral CB1 receptor antagonist JD5037 attenuates liver fibrosis via a CB1 receptor/β-arrestin1/Akt pathway, Br J Pharmacol 177 (2020) 2830–2847. 10.1111/bph.15010.

[25] M.E. Cooper, P.K. Nørregaard, T. Högberg, G. Andersson, J.-M. Receveur, J.-M. Linget, C.E. Elling, Efficacy in diet-induced obese mice of the hepatotropic, peripheral cannabinoid 1 receptor inverse agonist TM38837, Br J Pharmacol 181 (2024) 3926–3943. 10.1111/bph.16401.

[26] L. Maccioni, S. Dvorácskó, G. Godlewski, R. Cinar, M.R. Iyer, B. Gao, G. Kunos, Gut cannabinoid receptor 1 regulates alcohol binge-induced intestinal permeability, eGastroenterology 3 (2025) e100173. 10.1136/egastro-2024-100173.

[27] P.A. Perez, M.B. Wiley, A. Makriyannis, N.V. DiPatrizio, Cannabinoids Block Fat-induced Incretin Release via CB1-dependent and CB1-independent Pathways in Intestinal Epithelium, Gastro Hep Adv 3 (2024) 931–941. 10.1016/j.gastha.2024.07.006.

[28] I. Ruiz de Azua, G. Mancini, R.K. Srivastava, A.A. Rey, P. Cardinal, L. Tedesco, C.M. Zingaretti, A. Sassmann, C. Quarta, C. Schwitter, A. Conrad, N. Wettschureck, V.K. Vemuri, A. Makriyannis, J. Hartwig, M. Mendez-Lago, L. Bindila, K. Monory, A. Giordano, S. Cinti, G. Marsicano, S. Offermanns, E. Nisoli, U. Pagotto, D. Cota, B. Lutz, Adipocyte cannabinoid receptor CB1 regulates energy homeostasis and alternatively activated macrophages, J Clin Invest 127 (2017) 4148–4162. 10.1172/JCI83626.

[29] S. Wang, Q. Zhu, G. Liang, T. Franks, M. Boucher, K.K. Bence, M. Lu, C.M. Castorena, S. Zhao, J.K. Elmquist, P.E. Scherer, J.D. Horton, Cannabinoid receptor 1 signaling in hepatocytes and stellate cells does not contribute to NAFLD, J Clin Invest 131 (2021) e152242. 10.1172/JCI152242.

[30] C.R. Vianna, J. Donato, J. Rossi, M. Scott, K. Economides, L. Gautron, S. Pierpont, C.F. Elias, J.K. Elmquist, Cannabinoid receptor 1 in the vagus nerve is dispensable for body weight homeostasis but required for normal gastrointestinal motility, J Neurosci 32 (2012) 10331–10337. 10.1523/JNEUROSCI.4507-11.2012.

[31] D.A. Argueta, P.A. Perez, A. Makriyannis, N.V. DiPatrizio, Cannabinoid CB1 Receptors Inhibit Gut-Brain Satiation Signaling in Diet-Induced Obesity, Front Physiol 10 (2019) 704. 10.3389/fphys.2019.00704.

[32] A.G. Sykaras, C. Demenis, R.M. Case, J.T. McLaughlin, C.P. Smith, Duodenal enteroendocrine I-cells contain mRNA transcripts encoding key endocannabinoid and fatty acid receptors, PLoS One 7 (2012) e42373. 10.1371/journal.pone.0042373.

[33] K.L. Egerod, N. Petersen, P.N. Timshel, J.C. Rekling, Y. Wang, Q. Liu, T.W. Schwartz, L. Gautron, Profiling of G protein-coupled receptors in vagal afferents reveals novel gut-to-brain sensing mechanisms, Mol Metab 12 (2018) 62–75. 10.1016/j.molmet.2018.03.016.

[34] G. Burdyga, S. Lal, A. Varro, R. Dimaline, D.G. Thompson, G.J. Dockray, Expression of cannabinoid CB1 receptors by vagal afferent neurons is inhibited by cholecystokinin, J Neurosci 24 (2004) 2708–2715. 10.1523/JNEUROSCI.5404-03.2004.

[35] N. Percie du Sert, V. Hurst, A. Ahluwalia, S. Alam, M.T. Avey, M. Baker, W.J. Browne, A. Clark, I.C. Cuthill, U. Dirnagl, M. Emerson, P. Garner, S.T. Holgate, D.W. Howells, N.A. Karp, S.E. Lazic, K. Lidster, C.J. MacCallum, M. Macleod, E.J. Pearl, O.H. Petersen, F. Rawle, P. Reynolds, K. Rooney, E.S. Sena, S.D. Silberberg, T. Steckler, H. Würbel, The ARRIVE guidelines 2.0: Updated guidelines for reporting animal research, PLoS Biol 18 (2020) e3000410. 10.1371/journal.pbio.3000410.

[36] E. Lilley, S.C. Stanford, D.E. Kendall, S.P.H. Alexander, G. Cirino, J.R. Docherty, C.H. George, P.A. Insel, A.A. Izzo, Y. Ji, R.A. Panettieri, C.G. Sobey, B. Stefanska, G. Stephens, M. Teixeira, A. Ahluwalia, ARRIVE 2.0 and the British Journal of Pharmacology: Updated guidance for 2020, Br J Pharmacol 177 (2020) 3611–3616. 10.1111/bph.15178.

[37] O. Onimus, F. Arrivet, I.N. de O. Souza, B. Bertrand, J. Castel, S. Luquet, J.-P. Mothet, N. Heck, G. Gangarossa, The gut-brain vagal axis scales hippocampal memory processes and plasticity, Neurobiol Dis 199 (2024) 106569. 10.1016/j.nbd.2024.106569.

[38] O. Onimus, F. Arrivet, T. Le Borgne, S. Perez, J. Castel, A. Ansoult, B. Bertrand, N. Mashhour, C. de Almeida, L.-C. Bui, M. Vandecasteele, S. Luquet, L. Venance, N. Heck, F. Marti, G. Gangarossa, The gut-brain vagal axis governs mesolimbic dopamine dynamics and reward events, Sci Adv 12 (2026) eadz0828. 10.1126/sciadv.adz0828.

[39] S. Udi, L. Hinden, M. Ahmad, A. Drori, M.R. Iyer, R. Cinar, M. Herman-Edelstein, J. Tam, Dual inhibition of cannabinoid CB1 receptor and inducible NOS attenuates obesity-induced chronic kidney disease, Br J Pharmacol 177 (2020) 110–127. 10.1111/bph.14849.

[40] P. Zizzari, R. He, S. Falk, L. Bellocchio, C. Allard, S. Clark, T. Lesté-Lasserre, G. Marsicano, C. Clemmensen, D. Perez-Tilve, B. Finan, D. Cota, C. Quarta, CB1 and GLP-1 Receptors Cross Talk Provides New Therapies for Obesity, Diabetes 70 (2021) 415–422. 10.2337/db20-0162.

[41] H. Ma, G. Zhang, C. Mou, X. Fu, Y. Chen, Peripheral CB1 Receptor Neutral Antagonist, AM6545, Ameliorates Hypometabolic Obesity and Improves Adipokine Secretion in Monosodium Glutamate Induced Obese Mice, Front Pharmacol 9 (2018) 156. 10.3389/fphar.2018.00156.

[42] D. Pérez-Tilve, L. González-Matías, B.A. Aulinger, M. Alvarez-Crespo, M. Gil-Lozano, E. Alvarez, A.M. Andrade-Olivie, M.H. Tschöp, D.A. D’Alessio, F. Mallo, Exendin-4 increases blood glucose levels acutely in rats by activation of the sympathetic nervous system, Am J Physiol Endocrinol Metab 298 (2010) E1088–1096. 10.1152/ajpendo.00464.2009.

[43] T.H. Claus, C.Q. Pan, J.M. Buxton, L. Yang, J.C. Reynolds, N. Barucci, M. Burns, A.A. Ortiz, S. Roczniak, J.N. Livingston, K.B. Clairmont, J.P. Whelan, Dual-acting peptide with prolonged glucagon-like peptide-1 receptor agonist and glucagon receptor antagonist activity for the treatment of type 2 diabetes, J Endocrinol 192 (2007) 371–380. 10.1677/JOE-06-0018.

[44] F.H.M. Do Monte, N.S. Canteras, D. Fernandes, J. Assreuy, A.P. Carobrez, New perspectives on beta-adrenergic mediation of innate and learned fear responses to predator odor, J Neurosci 28 (2008) 13296–13302. 10.1523/JNEUROSCI.2843-08.2008.

[45] L. Bellocchio, E. Soria-Gómez, C. Quarta, M. Metna-Laurent, P. Cardinal, E. Binder, A. Cannich, A. Delamarre, M. Häring, M. Martín-Fontecha, D. Vega, T. Leste-Lasserre, D. Bartsch, K. Monory, B. Lutz, F. Chaouloff, U. Pagotto, M. Guzman, D. Cota, G. Marsicano, Activation of the sympathetic nervous system mediates hypophagic and anxiety-like effects of CB[ receptor blockade, Proc Natl Acad Sci U S A 110 (2013) 4786–4791. 10.1073/pnas.1218573110.

[46] M.E. Olson, D. Vizzutti, D.W. Morck, A.K. Cox, The parasympatholytic effects of atropine sulfate and glycopyrrolate in rats and rabbits, Can J Vet Res 58 (1994) 254–258.

[47] G. Gangarossa, L. Castell, L. Castro, P. Tarot, F. Veyrunes, P. Vincent, F. Bertaso, E. Valjent, Contrasting patterns of ERK activation in the tail of the striatum in response to aversive and rewarding signals, J. Neurochem. (2019). 10.1111/jnc.14804.

[48] O. Onimus, E. Valjent, G. Fisone, G. Gangarossa, Haloperidol-Induced Immediate Early Genes in Striatopallidal Neurons Requires the Converging Action of cAMP/PKA/DARPP-32 and mTOR Pathways, Int J Mol Sci 23 (2022) 11637. 10.3390/ijms231911637.

[49] J. Castel, G. Li, O. Onimus, E. Leishman, P.D. Cani, H. Bradshaw, K. Mackie, A. Everard, S. Luquet, G. Gangarossa, NAPE-PLD in the ventral tegmental area regulates reward events, feeding and energy homeostasis, Mol Psychiatry 29 (2024) 1478–1490. 10.1038/s41380-024-02427-6.

[50] L. Bai, S. Mesgarzadeh, K.S. Ramesh, E.L. Huey, Y. Liu, L.A. Gray, T.J. Aitken, Y. Chen, L.R. Beutler, J.S. Ahn, L. Madisen, H. Zeng, M.A. Krasnow, Z.A. Knight, Genetic Identification of Vagal Sensory Neurons That Control Feeding, Cell 179 (2019) 1129–1143.e23. 10.1016/j.cell.2019.10.031.

[51] S.P.H. Alexander, E. Kelly, A.A. Mathie, J.A. Peters, E.L. Veale, J.F. Armstrong, O.P. Buneman, E. Faccenda, S.D. Harding, M. Spedding, J.A. Cidlowski, D. Fabbro, A.P. Davenport, J. Striessnig, J.A. Davies, K.E. Ahlers-Dannen, M. Alqinyah, T.V. Arumugam, C. Bodle, J.B. Dagner, B. Chakravarti, S.P. Choudhuri, K.M. Druey, R.A. Fisher, K.J. Gerber, J.R. Hepler, S.B. Hooks, H.S. Kantheti, B. Karaj, S. Layeghi-Ghalehsoukhteh, J.-K. Lee, Z. Luo, K. Martemyanov, L.D. Mascarenhas, H. McNabb, C. Montañez-Miranda, O. Ogujiofor, H. Phan, D.L. Roman, V. Shaw, B. Sjogren, C. Sobey, M.M. Spicer, K.E. Squires, L. Sutton, M. Wendimu, T. Wilkie, K. Xie, Q. Zhang, Y. Zolghadri, The Concise Guide to PHARMACOLOGY 2023/24: Introduction and Other Protein Targets, Br J Pharmacol 180 Suppl 2 (2023) S1–S22. 10.1111/bph.16176.

[52] J. Tam, L. Hinden, A. Drori, S. Udi, S. Azar, S. Baraghithy, The therapeutic potential of targeting the peripheral endocannabinoid/CB1 receptor system, Eur J Intern Med 49 (2018) 23–29. 10.1016/j.ejim.2018.01.009.

[53] I.C. Alcantara, A.P.M. Tapia, Y. Aponte, M.J. Krashes, Acts of appetite: neural circuits governing the appetitive, consummatory, and terminating phases of feeding, Nat Metab 4 (2022) 836–847. 10.1038/s42255-022-00611-y.

[54] G. D’Agostino, D. Lyons, C. Cristiano, M. Lettieri, C. Olarte-Sanchez, L.K. Burke, M. Greenwald-Yarnell, C. Cansell, B. Doslikova, T. Georgescu, P.B. Martinez de Morentin, M.G. Myers, J.J. Rochford, L.K. Heisler, Nucleus of the Solitary Tract Serotonin 5-HT2C Receptors Modulate Food Intake, Cell Metab 28 (2018) 619–630.e5. 10.1016/j.cmet.2018.07.017.

[55] M.R. Hayes, K.P. Skibicka, T.M. Leichner, D.J. Guarnieri, R.J. DiLeone, K.K. Bence, H.J. Grill, Endogenous leptin signaling in the caudal nucleus tractus solitarius and area postrema is required for energy balance regulation, Cell Metab 11 (2010) 77–83. 10.1016/j.cmet.2009.10.009.

[56] C.W. Roman, V.A. Derkach, R.D. Palmiter, Genetically and functionally defined NTS to PBN brain circuits mediating anorexia, Nat Commun 7 (2016) 11905. 10.1038/ncomms11905.

[57] S. Engeli, J. Böhnke, M. Feldpausch, K. Gorzelniak, J. Janke, S. Bátkai, P. Pacher, J. Harvey-White, F.C. Luft, A.M. Sharma, J. Jordan, Activation of the peripheral endocannabinoid system in human obesity, Diabetes 54 (2005) 2838–2843. 10.2337/diabetes.54.10.2838.

[58] I. Matias, B. Gatta-Cherifi, A. Tabarin, S. Clark, T. Leste-Lasserre, G. Marsicano, P.V. Piazza, D. Cota, Endocannabinoids measurement in human saliva as potential biomarker of obesity, PLoS One 7 (2012) e42399. 10.1371/journal.pone.0042399.

[59] N. Mattelaer, B. Van der Schueren, L. Van Oudenhove, N. Weltens, R. Vangoitsenhoven, The circulating and central endocannabinoid system in obesity and weight loss, Int J Obes (Lond) 48 (2024) 1363–1382. 10.1038/s41366-024-01553-z.

[60] I. González-Mariscal, S.M. Krzysik-Walker, W. Kim, M. Rouse, J.M. Egan, Blockade of cannabinoid 1 receptor improves GLP-1R mediated insulin secretion in mice, Mol Cell Endocrinol 423 (2016) 1–10. 10.1016/j.mce.2015.12.015.

[61] C.E. Moss, W.J. Marsh, H.E. Parker, E. Ogunnowo-Bada, C.H. Riches, A.M. Habib, M.L. Evans, F.M. Gribble, F. Reimann, Somatostatin receptor 5 and cannabinoid receptor 1 activation inhibit secretion of glucose-dependent insulinotropic polypeptide from intestinal K cells in rodents, Diabetologia 55 (2012) 3094–3103. 10.1007/s00125-012-2663-5.

[62] H.-R. Berthoud, V.L. Albaugh, W.L. Neuhuber, Gut-brain communication and obesity: understanding functions of the vagus nerve, J Clin Invest 131 (2021) e143770, 143770. 10.1172/JCI143770.

[63] C. Clemmensen, T.D. Müller, S.C. Woods, H.-R. Berthoud, R.J. Seeley, M.H. Tschöp, Gut-Brain Cross-Talk in Metabolic Control, Cell 168 (2017) 758–774. 10.1016/j.cell.2017.01.025.

[64] J.C. Brüning, H. Fenselau, Integrative neurocircuits that control metabolism and food intake, Science 381 (2023) eabl7398. 10.1126/science.abl7398.

[65] C. Quarta, L. Bellocchio, G. Mancini, R. Mazza, C. Cervino, L.J. Braulke, C. Fekete, R. Latorre, C. Nanni, M. Bucci, L.E. Clemens, G. Heldmaier, M. Watanabe, T. Leste-Lassere, M. Maitre, L. Tedesco, F. Fanelli, S. Reuss, S. Klaus, R.K. Srivastava, K. Monory, A. Valerio, A. Grandis, R. De Giorgio, R. Pasquali, E. Nisoli, D. Cota, B. Lutz, G. Marsicano, U. Pagotto, CB(1) signaling in forebrain and sympathetic neurons is a key determinant of endocannabinoid actions on energy balance, Cell Metab 11 (2010) 273–285. 10.1016/j.cmet.2010.02.015.

[66] M. Côté, I. Matias, I. Lemieux, S. Petrosino, N. Alméras, J.-P. Després, V. Di Marzo, Circulating endocannabinoid levels, abdominal adiposity and related cardiometabolic risk factors in obese men, Int J Obes (Lond) 31 (2007) 692–699. 10.1038/sj.ijo.0803539.

[67] I. Matias, S. Petrosino, A. Racioppi, R. Capasso, A.A. Izzo, V. Di Marzo, Dysregulation of peripheral endocannabinoid levels in hyperglycemia and obesity: Effect of high fat diets, Mol Cell Endocrinol 286 (2008) S66–78. 10.1016/j.mce.2008.01.026.

[68] D.M. Daly, S.J. Park, W.C. Valinsky, M.J. Beyak, Impaired intestinal afferent nerve satiety signalling and vagal afferent excitability in diet induced obesity in the mouse, J Physiol 589 (2011) 2857–2870. 10.1113/jphysiol.2010.204594.

[69] G. de Lartigue, Role of the vagus nerve in the development and treatment of diet-induced obesity, J Physiol 594 (2016) 5791–5815. 10.1113/JP271538.

[70] G. de Lartigue, C. Barbier de la Serre, E. Espero, J. Lee, H.E. Raybould, Diet-induced obesity leads to the development of leptin resistance in vagal afferent neurons, Am J Physiol Endocrinol Metab 301 (2011) E187–195. 10.1152/ajpendo.00056.2011.

[71] H. Loper, M. Leinen, L. Bassoff, J. Sample, M. Romero-Ortega, K.J. Gustafson, D.M. Taylor, M.A. Schiefer, Both high fat and high carbohydrate diets impair vagus nerve signaling of satiety, Sci Rep 11 (2021) 10394. 10.1038/s41598-021-89465-0.

[72] G. Godlewski, R. Cinar, N.J. Coffey, J. Liu, T. Jourdan, B. Mukhopadhyay, L. Chedester, Z. Liu, D. Osei-Hyiaman, M.R. Iyer, J.K. Park, R.G. Smith, H. Iwakura, G. Kunos, Targeting Peripheral CB1 Receptors Reduces Ethanol Intake via a Gut-Brain Axis, Cell Metab 29 (2019) 1320–1333.e8. 10.1016/j.cmet.2019.04.012.

[73] A. Herrerias, A. Oliverio, S. Dvorácskó, A. Thyagarajan, L. Chedester, J. Liu, R. Cinar, M.R. Iyer, G. Kunos, G. Godlewski, CB1 receptors on a subset of vagal afferent neurons modulate voluntary ethanol intake in mice, Mol Psychiatry (2025). 10.1038/s41380-025-03266-9.

[74] R. Gómez, M. Navarro, B. Ferrer, J.M. Trigo, A. Bilbao, I. Del Arco, A. Cippitelli, F. Nava, D. Piomelli, F. Rodríguez de Fonseca, A peripheral mechanism for CB1 cannabinoid receptor-dependent modulation of feeding, J Neurosci 22 (2002) 9612–9617.

[75] C.P. Wood, C. Alvarez, N.V. DiPatrizio, Cholinergic Neurotransmission Controls Orexigenic Endocannabinoid Signaling in the Gut in Diet-Induced Obesity, J Neurosci 44 (2024) e0813232024. 10.1523/JNEUROSCI.0813-23.2024.

[76] N.P. Bowles, I.N. Karatsoreos, X. Li, V.K. Vemuri, J.-A. Wood, Z. Li, K.L.K. Tamashiro, G.J. Schwartz, A.M. Makriyannis, G. Kunos, C.J. Hillard, B.S. McEwen, M.N. Hill, A peripheral endocannabinoid mechanism contributes to glucocorticoid-mediated metabolic syndrome, Proc Natl Acad Sci U S A 112 (2015) 285–290. 10.1073/pnas.1421420112.

[77] A. Drori, A. Gammal, S. Azar, L. Hinden, R. Hadar, D. Wesley, A. Nemirovski, G. Szanda, M. Salton, B. Tirosh, J. Tam, CB1R regulates soluble leptin receptor levels via CHOP, contributing to hepatic leptin resistance, Elife 9 (2020) e60771. 10.7554/eLife.60771.

[78] A. Ghosh, M.-L. Peyot, Y.H. Leung, F. Ravenelle, S.R.M. Madiraju, M. Prentki, A peripherally restricted cannabinoid-1 receptor inverse agonist promotes insulin secretion and protects from cytokine toxicity in human pancreatic islets, Eur J Pharmacol 944 (2023) 175589. 10.1016/j.ejphar.2023.175589.

[79] T. Jourdan, J.K. Park, Z.V. Varga, J. Pálóczi, N.J. Coffey, A.Z. Rosenberg, G. Godlewski, R. Cinar, K. Mackie, P. Pacher, G. Kunos, Cannabinoid-1 receptor deletion in podocytes mitigates both glomerular and tubular dysfunction in a mouse model of diabetic nephropathy, Diabetes Obes Metab 20 (2018) 698–708. 10.1111/dom.13150.

[80] H. Münzberg, H.-R. Berthoud, W.L. Neuhuber, Sensory spinal interoceptive pathways and energy balance regulation, Molecular Metabolism 78 (2023) 101817. 10.1016/j.molmet.2023.101817.

[81] G. Veress, Z. Meszar, D. Muszil, A. Avelino, K. Matesz, K. Mackie, I. Nagy, Characterisation of cannabinoid 1 receptor expression in the perikarya, and peripheral and spinal processes of primary sensory neurons, Brain Struct Funct 218 (2013) 733–750. 10.1007/s00429-012-0425-2.

[82] V. Rakotoarivelo, B. Allam-Ndoul, C. Martin, L. Biertho, V. Di Marzo, N. Flamand, A. Veilleux, Investigating the alterations of endocannabinoidome signaling in the human small intestine in the context of obesity and type 2 diabetes, Heliyon 10 (2024) e26968. 10.1016/j.heliyon.2024.e26968.

[83] J.A. Noriega-Prieto, R. Falcón-Moya, J.A. Noeker, R. Cai, U.B. Fundazuri, A. Eraso-Pichot, S. Cai, P. Guttipatti, L. Belisle, A. Rodríguez-Moreno, M. van der Stelt, Y. Li, J.F. Cheer, G. Marsicano, P. Kofuji, A. Araque, Distinct endocannabinoids specifically signal to astrocytes or neurons in the adult mouse hippocampus, Nat Neurosci 29 (2026) 445–454. 10.1038/s41593-025-02148-1.

[84] W. Han, L.A. Tellez, M.H. Perkins, I.O. Perez, T. Qu, J. Ferreira, T.L. Ferreira, D. Quinn, Z.-W. Liu, X.-B. Gao, M.M. Kaelberer, D.V. Bohórquez, S.J. Shammah-Lagnado, G. de Lartigue, I.E. de Araujo, A Neural Circuit for Gut-Induced Reward, Cell 175 (2018) 665–678.e23. 10.1016/j.cell.2018.08.049.

[85] A. Bergadà-Martínez, L. de Los Reyes-Ramírez, S. Martínez-Torres, L. Ciaran-Alfano, I. Martínez-Gallego, R. Maldonado, A. Rodríguez-Moreno, A. Ozaita, Sub-chronic administration of AM6545 enhances cognitive performance and induces hippocampal synaptic plasticity changes in naïve mice, Br J Pharmacol (2025). 10.1111/bph.70015.

[86] A. Busquets-Garcia, M. Gomis-González, R.K. Srivastava, L. Cutando, A. Ortega-Alvaro, S. Ruehle, F. Remmers, L. Bindila, L. Bellocchio, G. Marsicano, B. Lutz, R. Maldonado, A. Ozaita, Peripheral and central CB1 cannabinoid receptors control stress-induced impairment of memory consolidation, Proc Natl Acad Sci U S A 113 (2016) 9904–9909. 10.1073/pnas.1525066113.

[87] S. Martínez-Torres, A. Bergadà-Martínez, J.E. Ortega, L. Galera-López, A. Hervera, L. de Los Reyes-Ramírez, A. Ortega-Álvaro, F. Remmers, E. Muñoz-Moreno, G. Soria, J.A. Del Río, B. Lutz, J.Á. Ruíz-Ortega, J.J. Meana, R. Maldonado, A. Ozaita, Peripheral CB1 receptor blockade acts as a memory enhancer through a noradrenergic mechanism, Neuropsychopharmacology 48 (2023) 341–350. 10.1038/s41386-022-01436-9.

[88] F.K. Knop, G. Kunos, D. Dicker, J.-S. Paquette, L. Aronne, O. Frenkel, T. Holst-Hansen, K. Lalonde, J. Lee, G. Crater, trial investigators, Efficacy and safety of monlunabant in adults with obesity and metabolic syndrome: a double-blind, randomised, placebo-controlled, phase 2a trial, Lancet Diabetes Endocrinol 13 (2025) 911–923. 10.1016/S2213-8587(25)00216-5.

[89] G.D. Crater, K. Lalonde, F. Ravenelle, M. Harvey, J.-P. Després, Effects of CB1R inverse agonist, INV-202, in patients with features of metabolic syndrome. A randomized, placebo-controlled, double-blind phase 1b study, Diabetes Obes Metab 26 (2024) 642–649. 10.1111/dom.15353.

